# The Histone Chaperone Spn1 Preserves Chromatin Protections at Promoters and Nucleosome Positioning in Open Reading Frames

**DOI:** 10.1101/2024.03.14.585010

**Authors:** Andrew J. Tonsager, Alexis Zukowski, Catherine A. Radebaugh, Abigail Weirich, Laurie A. Stargell, Srinivas Ramachandran

## Abstract

Spn1 is a multifunctional histone chaperone that associates with RNA polymerase II during elongation and is essential for life in eukaryotes. While previous work has elucidated regions of the protein important for its many interactions, it is unknown how these domains contribute to the maintenance of chromatin structure. Here, we employ digestion by micrococcal nuclease followed by single-stranded library preparation and sequencing (MNase-SSP) to characterize chromatin structure in *Saccharomyces cerevisiae* expressing wild-type or mutants of Spn1 (*spn1^K192N^ or spn1^141-305^*). We mapped protections of all sizes genome-wide. Surprisingly, we observed a widespread loss of short fragments over nucleosome-depleted regions (NDRs) at promoters in the *spn1^K192N^*-containing strain, indicating critical functions of Spn1 in maintaining normal chromatin architecture outside open reading frames. Additionally, there are shifts in DNA protections in both Spn1 mutant expressing strains over open reading frames, which indicate changes in nucleosome and subnucleosome positioning. This was observed in markedly different Spn1 mutant strains, demonstrating that multiple functions of Spn1 are required to maintain proper chromatin structure in open reading frames. Changes in chromatin structure correlate positively with changes in gene expression as shown by RNA-seq analysis in the Spn1 mutant strains. Taken together, our results reveal a previously unknown role of Spn1 in the maintenance of NDR architecture and deepen our understanding of Spn1-dependent chromatin maintenance over transcribed regions.

## INTRODUCTION

Eukaryotes package their genomes by wrapping ∼147 bp of DNA in about 1.7 superhelical turns around an octamer of histone proteins to form nucleosomes (Davey et al. 2002; Flaus et al. 1996; Luger et al. 1997). This process occurs in a stepwise manner, as DNA initially wraps around a tetramer of histones H3–H4 to form a tetrasome, followed by sequential additions of two H2A-H2B dimers to form a hexasome then a nucleosome (Polo and Almouzni 2006). Tetrasome and hexasome subnucleosomal structures have been observed both *in vitro* (Brower-Toland et al. 2002) and *in vivo* (Ramachandran et al. 2017) and represent stable subnucleosomal intermediates in nucleosome assembly and disassembly. Nucleosomes regulate DNA access to many proteins that perform essential processes such as DNA replication (Eaton et al. 2010; Groth et al. 2007), repair (Adkins et al. 2013; Groth et al. 2007), recombination (Bevington and Boyes 2013), and transcription. Nucleosomes regulate transcription in a variety of ways. Positioning of nucleosomes at promoters can hinder transcription initiation by preventing the binding of transcription factors (Henikoff et al. 2011; Kubik et al. 2015) and nucleosome-depleted regions (NDRs) are important for regulation of gene expression (Kubik et al. 2015). Chromatin remodelers, such as the RSC complex, actively maintain NDRs after nucleosome deposition (Lorch et al. 2014). This region is bound by many other factors, such as activators that bind and compete with nucleosome formation within NDRs, contributing to the low nucleosome density in these regions (Rando and Winston 2012). NDRs upstream of transcription start sites contain promoter sequences bound by transcription initiation factors. Positioning of nucleosomes within coding regions can play an important role in preventing intragenic transcription initiation by reducing the accessibility of factors to intragenic initiation sites (Kaplan et al. 2003). Nucleosomes inhibit transcription elongation as they pose a significant barrier to RNA polymerase II (RNAPII) and reduce its transcription rate (Brahma and Henikoff 2020; Selth et al. 2010). Hence, chromatin structure *in vivo* must be dynamic to allow for selective gene expression. An important class of proteins responsible for modifying chromatin structure are histone chaperones, which bind histones and facilitate histone deposition, exchange, or removal from chromatin, independent of ATP (Bai and Morozov 2010).

Spn1 is a multifunctional and essential histone chaperone that is highly conserved across eukaryotes. *S. cerevisiae* Spn1 preferentially binds H3-H4 histones (Li et al. 2018) via its N-terminal region, with a high binding affinity (K_D_ ∼ 10nM). Spn1 affinity to H3-H4 is at least 50-fold higher than that to H2A-H2B dimers (Li et al. 2022). Spn1 can deposit H3-H4 onto DNA and facilitate the assembly of H3-H4 tetrasomes *in vitro* (Li et al. 2018). Spn1 also has a well-characterized interaction with Spt6, another essential histone chaperone; they bind each other with high affinity (K_D_ = 10-100nM) to form a heterodimer (Gavin et al. 2002; Krogan et al. 2002; Lindstrom et al. 2003; McDonald et al. 2010). The central domain of Spn1 is a highly conserved eight-helix bundle and is primarily responsible for binding Spt6 (Li et al. 2022). Depletion of either protein through the auxin-inducible degron system results in a reduction in recruitment of the other to chromatin (Reim et al. 2020). Spn1 binds nucleosomes with its C-terminal region (Li et al. 2018; Li et al. 2022).

A mutant of Spn1, *spn1^141-305^*, lacks the N-and C-terminal domains, which are responsible for histone and nucleosome binding, respectively. When *spn1^141-305^* is the sole source of Spn1, yeast cell growth is relatively healthy (Fischbeck et al. 2002; Li et al. 2018). Though *spn1^141-305^*cannot bind histones and nucleosomes, it retains the ability to bind Spt6 and RNAPII (Li et al. 2018; Li et al. 2022). A different allele of Spn1, *spn1^K192N^,* is similarly viable, but we have previously demonstrated that these cells have reduced Spn1 recruitment to the *CYC1* locus and fail to recruit Spt6 to *CYC1* (Zhang et al. 2008). The *spn1^K192N^* mutant also has reduced Spn1 binding to RNAPII by co-immunoprecipitation (Zhang et al. 2008). While both alleles cover a *SPN1* knockout, we have previously shown that they are partially defective or involved in redundant functions with other cellular proteins. For example, yeast expressing the Spn1 mutants in a background containing knockouts in other elongation and transcription factors experience significant defects in growth (Li et al. 2018; Zhang et al. 2008). Recent work demonstrates that Iws1, the human ortholog of Spn1, serves as a hub that maintains association of numerous additional transcription factors (Cermakova et al. 2021).

While we and others have previously demonstrated that Spn1 possesses multiple interactions pertinent to its chromatin functions, the role of Spn1 *in vivo* is not well understood. Here, micrococcal nuclease digestion followed by single-stranded library preparation and sequencing (MNase-SSP) (Ramani et al. 2019) is used to characterize chromatin structure in yeast expressing wild-type or mutant Spn1 alleles. Compared to traditional MNase-seq (with or without selection of ∼147 bp fragments), MNase-SSP allows for the analysis of a wider range of protections by better retaining MNase-protected fragments of a variety of lengths (Ramani et al. 2019). This includes both nucleosomal protections and subnucleosomal protections. Subnucleosome protections are broadly defined as DNA protections with footprints smaller than the protection by a nucleosome (∼147bp) (Henikoff et al. 2011; Kent et al. 2011). Specific subnucleosome protections can arise due to histone-containing subnucleosomal intermediates at open reading frames (Ramachandran et al. 2017; Rhee et al. 2014). The MNase-SSP protections also include non-nucleosomal protections, defined as DNA fragments obtained through non-histone protections, such as protection by transcription factors, pre-initiation complex, and other DNA binding factors. In this study, using MNase-SSP, we outline the effects of *spn1^K192N^* and *spn1^141-305^* on chromatin structure *in vivo*. Furthermore, to determine whether changes in chromatin structure identified through MNase-SSP correlate with changes in gene expression, we also performed RNA-seq to identify differentially expressed genes in cells expressing *spn1^K192N^*or *spn1^141-305^*.

We find that despite distinct defects with known binding partners, both *spn1^K192N^* and *spn1^141-305^* expression show similar chromatin defects across genes, namely a progressive 3’ downstream shift in chromatin protections over gene bodies. Intriguingly, MNase-SSP also revealed a molecular mutant phenotype specific to cells expressing *spn1^K192N^*: the loss of protections over nucleosome-depleted regions (NDRs) in promoters, most dramatic for fragments <60bp in length. Additionally, expression of *spn1^141-305^*, but not *spn1^K192N^*, produces a unique molecular mutant phenotype of a shift in the +1 and +2 nucleosome positions. Mapping the midpoints of the sequenced fragments indicates that shifts correspond to a downstream redistribution of nucleosome and subnucleosomal occupancy at alternative rotational positions; alternative rotational positions have been previously observed in the yeast genome (Brogaard et al. 2012). This molecular mutant phenotype of the shift downstream is largest at longer genes. Our RNA-seq results also revealed that expression of *spn1^K192N^* or *spn1^141-305^* produces global reduction in gene expression. Intriguingly, genes differentially expressed in the Spn1 mutant strains are enriched in longer genes, revealing a correlative relationship between instances of nucleosome positioning shift and changes in gene expression. Taken together, these studies reveal the importance of Spn1 in preserving chromatin structure across the yeast genome.

## MATERIAL AND METHODS

### Yeast Culturing and Strains

All *Saccharomyces cerevisiae* strains were grown in yeast extract, peptone, and dextrose (YPD; 2% dextrose) media at 30°C in biological duplicate independently. A description of yeast strains and plasmids is provided in Supplemental Table S1. The strains used in this study were previously created as described (Li et al. 2018; Thurston et al. 2018; Zhang et al. 2008). In summary, strains were transformed with a covering plasmid containing the *URA3* gene and the wild-type *SPN1* open reading frame flanked by the *TOA1* promoter and terminator; the *TOA1* sequences were used to prevent integration within the covering plasmid of the *LEU2* derivative described next. Endogenous *SPN1* was replaced by a *LEU2* fragment flanked by the *SPN1* promoter and terminator through homologous recombination. Plasmids containing the *HIS3* gene and wild-type *SPN1*, *spn1^K192N^*, or *spn1^141-305^* under the control of their native promoter and terminator were introduced into strains by plasmid shuffling, which results in removal of the covering plasmid.

### RNA preparation for sequencing

RNA was isolated from yeast expressing either wild-type Spn1, *spn1^K192N^*, or *spn1^141-305^* in triplicate 10-ml cultures by hot-acid phenol extraction as previously described (Iyer and Struhl 1996). Cells were grown in YPD at 30°C to mid-exponential phase (OD600 ∼ 1.0), harvested, and each sample was spiked to obtain a final concentration of 1 *S. pombe* to 10 *S. cerevisae* cells. RNA libraries were prepared with poly-A enrichment and sequenced using the NovaSeq X Plus Series (PE150) platform (Novogene).

### RNA-seq analysis

A concatenated fasta file was created using i) cDNA sequences of *S. cerevisiae* (release 113 from Ensembl), ii) cDNA sequences of *S. pombe* (from PomBase), iii) whole genome sequence of *S. cerevisiae*, and iv) whole genome sequence of *S. pombe*. A Salmon index was created using the concatenated fasta, setting the whole genome sequences as decoys. Transcript quantification was performed using Salmon v1.9 with libType set as automatic (Patro et al. 2017). Differential gene expression analysis was performed using DeSeq2 (Love et al. 2014) (v1.30.1) in R (v4.0.3). For spike-in normalization, size factors were estimated by setting *S. pombe* genes as control genes, resulting in the distribution of log2FC for S. pombe genes centered at 0 (**Figure S18A-C, File S1**). The expression matrix across genotypes was transformed using the “rlog” function in DESeq2 and the transformed matrix was used for performing principal component analysis (PCA) using the “prcomp” function in R and plotted using ggplot2. Gene set enrichment analysis was performed using g:Profiler (Kolberg et al. 2023).

### Isolation of nuclei and Micrococcal nuclease digestion

Yeast nuclei isolation and MNase digestion were performed as described in (Zhang and Reese 2005) with modifications. Yeast strains were grown in 400 ml of YP +2% glucose until an OD_600_ of ∼1.0 was reached. The cells were collected by centrifugation in a JA-10 rotor at 5,000 RPM for 10 minutes at 4°C. The supernatant was decanted from above the cell pellets, and the cells were suspended in 30 ml of Sorbitol buffer (50 mM Tris-Cl pH 7.5, 1.0 M Sorbitol, 10 mM MgCl_2_, 10 mM 2-Mercaptoethanol and 1mM Phenylmethylsulfonyl fluoride (PMSF)). The cell suspensions were transferred to chilled 50 ml conical tubes and collected by centrifugation in a GPKR centrifuge at 3,200 RPM for 5 minutes at 4°C. The supernatants were decanted from above the cell pellets and the cells were washed two additional times with 30 mL of Sorbitol buffer. The mass of the cell pellets was determined, and the cells were suspended in 0.66 ml of Sorbitol buffer per gram of cells.

Spheroplasting was completed by adding 0.34 mL of Zymolase (10 ^mg^/_mL_) per gram of cells and incubating at 30°C with gentle agitation until ∼90% of the cells were converted to spheroplasts. The spheroplasts were collected by centrifugation in a GPKR centrifuge at 3,200 RPM for 5 minutes at 4°C and then gently washed two times with 5 ml of Sorbitol buffer as described above. The washed spheroplasts were suspended in 25 mL of Ficoll buffer (18% Ficoll 400, 20 mM Tris-Cl pH 7.5, and 0.5 mM MgCl_2_), transferred to a 50 ml Potter-Elvehjem homogenizer and lysed on ice with six up-down even strokes with a handheld drill at max rpm. The lysates were transferred to chilled 25 ml centrifuge tubes, and the remaining cells were pelleted in a JA-20 rotor at 6400 x rpm for 5 minutes at 4°C. The supernatants containing yeast nuclei were transferred to new centrifuge tubes and pelleted at 15,000 x rpm at 4°C. The nuclei were suspended in 10 ml of MNase digestion buffer (20mM Tris-Cl pH 8.0, 1.5mM CaCl_2_, 150mM NaCl, and 1mM PMSF). To estimate the yield of nuclei, 100 µl aliquots of the nuclei suspensions were diluted to 1 ml with MNase digestion buffer and the OD_600_ was determined. The nuclei were again pelleted at 15,000 x rpm for 30 minutes at 4°C and suspended in MNase digestion buffer at a ratio of 2.4 ml per 0.2 OD_600_. Five aliquots (400 µl) of the nuclei suspensions were transferred to microcentrifuge tubes and incubated at 37°C for 10 minutes. Digestion buffer or MNase was added to samples to yield a final concentration of 0, 32, 64, or 128 U/ml, and incubation at 37°C continued for 10 minutes. 100 µl of stop solution (50mM EDTA, 2.5% SDS, 0.25mg/ml Proteinase K) was added to each reaction, and the samples were mixed and incubated at 37°C overnight. The MNase-digested DNA samples were extracted twice with phenol:chloroform, once with chloroform, and precipitated with ethanol. The DNA was washed with 70% ethanol, dried at room temperature, then suspended in 100 µl of TE buffer (pH 7.5) and 3 µl of RNaseA (10 mg/ml) added. The RNase reactions were incubated overnight at 37°C, followed by ethanol precipitation, washing with 70% ethanol, and drying at room temperature. The precipitated DNA samples were suspended in 100µl of TE buffer (10mM Tris-Cl, pH 8.0 and 1mM EDTA, pH 8.0), and 10µl of each sample was subjected to agarose gel electrophoresis.

### Single-stranded DNA library preparation (SSP) and sequencing

MNase-protected DNA fragments were made into sequencing libraries using the single-stranded library protocol as previously described (Gansauge and Meyer 2013; Ramani et al. 2019; Snyder et al. 2016). In brief, 10ng DNA was dephosphorylated using FastAP Thermosensitive Alkaline Phosphatase (Thermo Scientific cat. EF0651), denatured, and incubated overnight with CircLigaseII (Lucigen cat. CL9025K) and 0.093-0.125 µM biotinylated CL78 primer (35) at 60°C with shaking every 5 minutes. DNA fragments were then denatured and bound to magnetic streptavidin M-280 beads (Invitrogen cat. 11205D) for 30 minutes at room temperature with nutation. Beads were washed, and second-strand synthesis was performed using Bst 2.0 DNA polymerase (NEB cat. M0537) with an increasing temperature gradient of 15-31°C with shaking at 1750 rpm. Beads were washed, and a 3’ gap fill was performed using T4 DNA polymerase (Thermo Scientific cat. EL0011) for 30 minutes at room temperature. Beads were washed, and a double-stranded adapter was ligated using T4 DNA ligase (Thermo Scientific cat. EP0062) for 2 hours at room temperature with shaking at 1750 rpm. Beads were washed and resuspended in 30 µL 10 mM TET buffer (10 mM Tris-HCl pH 8.0, 1 mM EDTA pH 8.0, 0.05% Tween-20). DNA was denatured at 95°C for 3 min, and DNA libraries were collected after immediate magnetic separation.

Quantitative real-time PCR was performed on DNA libraries using iTAQ Supermix (Bio-Rad cat. 1725124), and Ct values were used to determine the number of PCR cycles needed to amplify each library. PCR was performed with KAPA HiFi DNA polymerase (Kapa Biosystems cat. KK2502) using barcoded indexing primers for Illumina. Primer dimers were removed from the libraries using AMPure beads (Beckman Coulter cat. A63881). Libraries were eluted in 0.1X TE and concentrations were determined using Qubit. The length distribution of each library was assessed by the Agilent Bioanalyzer using the D1000 or HSD1000 cassette. Libraries were sequenced for 150 cycles in paired-end mode on the NovaSeq 6000 system at the University of Colorado Cancer Center Genomics Shared Resource.

### MNase-SSP analysis

Sequencing reads were trimmed using Cutadapt (Martin 2011): Illumina adapter sequences were removed, and reads were trimmed to 140 bp. Reads less than 35 bp were discarded. Trimmed reads were aligned to the sacCer3 version of *Saccharomyces cerevisiae* genome using bowtie2 (Martin 2011). Samtools (Li et al. 2009) and bedtools (Quinlan and Hall 2010) were used for processing aligned reads from sam to bed files. Coverage at 1 bp windows genome-wide for specific fragment lengths was calculated as the number of reads that mapped at each window, normalized by the factor N: N = 139712364/(Total number of mapped reads). 139712364 was a number chosen arbitrarily. Coverage of fragment centers for specific fragment lengths was also calculated similarly at 1 bp windows. V-plots were generated using custom scripts that are publicly available (https://github.com/srinivasramachandran/VPLOTS/). Nucleosome positions were obtained from (Brogaard et al. 2012). Transcription start sites were obtained from (Park et al. 2014). Genes were stratified by expression using published NET-seq data (Churchman and Weissman 2011). NET-seq normalized read counts were summed between TSS+100 and TSS+200. Z-scores across rotational positions spanning −50 to +50 bp surrounding each nucleosome dyad position were calculated for each mapped dyad for each gene. Positive Z-scores denote preferred rotational positions within a nucleosome position of one gene. The Z-scores of wild-type Spn1 were subtracted from the Z-scores of Spn1 mutants for each gene to obtain changes in rotation position preference due to Spn1 mutation.

## RESULTS

### MNase-SSP interrogates chromatin protections across the yeast genome

To study the role of Spn1 in maintaining chromatin structure, wild-type Spn1, *spn1^K192N^*, and *spn1^141-305^*containing strains were subjected to MNase-SSP. MNase-SSP was performed with increasing levels of MNase, which digests linker DNA at 25 times the rate of nucleosomal DNA and produces protected genomic fragments of varied sizes (Chereji et al. 2017). Samples were subjected to agarose gel electrophoresis to qualitatively assess the population of polynucleosomal, nucleosomal, and smaller fragment sizes after MNase digestion (**Figure S1A, File S1**). Fragments of all lengths were sequenced and mapped to the *S. cerevisiae* genome. A metagene analysis was performed to generate plots of fragment midpoint versus fragment length (“V-plots”) (Henikoff et al. 2011) centered at the dyad of each nucleosome from the +1 to +10 positions. V-plots centered at +1, +2, and +5 exemplify that plots at each nucleosome position show a checkered pattern that appears similar across both wild-type (**Figure 1A**) and mutant samples (**Figure 1B, C**). To construct V-plots, we randomly sampled fragments genome-wide to yield similar fragment length distributions across WT and Spn1 mutant datasets (**Figure S1B, File S1**). Dyad definitions from chemical cleavage mapping were used to define nucleosome dyad positions spanning 4688 yeast genes (Brogaard et al. 2012) (**Figure S2, File S1**). The density at position 0 and fragment length 147 bp corresponds to the strongest dyad signal (**Figure 1**). For fragments ∼150 bp in length, this arises from peaks in density separated by a 10 bp periodicity on either side of the dyad, suggesting nucleosome occupancy at previously mapped rotational positions surrounding yeast nucleosomes (Brogaard et al. 2012; Chereji et al. 2018). Additionally, the V-plots indicate the existence of rotational positioning of subnucleosomal protections in all the samples, as this pattern persists at shorter fragment lengths. Notably, at ∼50-150 bp upstream of the +1 dyad, the *spn1^K192N^* mutation, but not *spn1^141-305^*, produces a loss in fragment density for short protections, corresponding to a decrease in protection over nucleosome-depleted regions (NDR) (**Figure 1B**). This finding suggests that Spn1 may contribute towards maintenance of the architecture of NDRs, a newly described function for Spn1.

**Figure 1.**
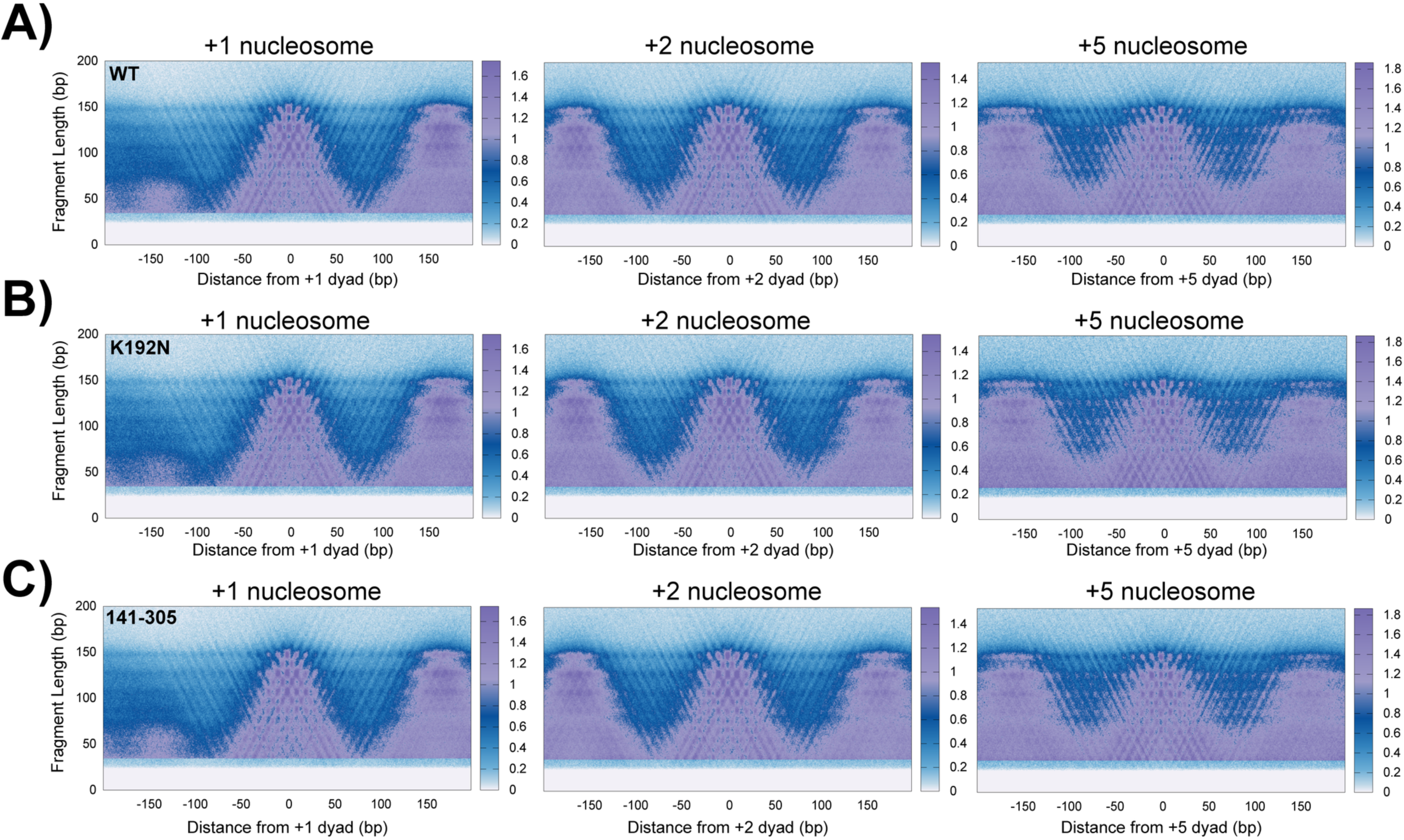
An extensive characterization of fragments protected from MNase digestion suggests shifts in chromatin protections in Spn1 mutant yeast across fragment lengths. (A) Nuclei harvested from two biological replicates of wild-type, (B) *spn1^K192N^*, and (C) *spn1^141-305^* expressing yeast grown to log phase were digested with 128U MNase and protected fragments were isolated and sequenced. V-plots were produced by mapping fragment midpoints to the yeast genome anchored at +1, +2, and +5 nucleosome dyad positions from 4688 genes.

### The *spn1^K192N^* mutation reduces stability of chromatin protections over nucleosome-depleted regions

The V-plot analyses indicate that MNase-protected fragments are reduced in the nucleosome-depleted regions (NDRs) specifically in the *spn1^K192N^* mutant yeast strain (**Figure 1**). NDRs are flanked by well-positioned −1 and +1 nucleosomes, which reside directly upstream and downstream of this region respectively. There are many contributors to the architecture of NDRs, including chromatin remodelers, activators, transcription initiation factors, etc. (Lorch et al. 2014; Rando and Winston 2012). Many of these non-nucleosomal factors have small footprints (<60 bp) (Henikoff et al. 2011; Kent et al. 2011). The V-plot analyses suggested that short protections of these lengths are reduced over the NDR in *spn1^K192N^* cells. To further explore this, we mapped the enrichment of short (<60 bp) fragments relative to the +1 nucleosome position of genes. This range of fragments produces a significant peak between the locations of the +1 and −1 nucleosomes in wild-type cells, and the abundance of fragments within this peak is dramatically reduced in *spn1^K192N^*, but not *spn1^141-305^* cells (**Figure 2A**). The enrichment of <60 bp fragments shows that short non-nucleosomal protections are lost over NDRs in *spn1^K192N^* cells.

**Figure 2.**
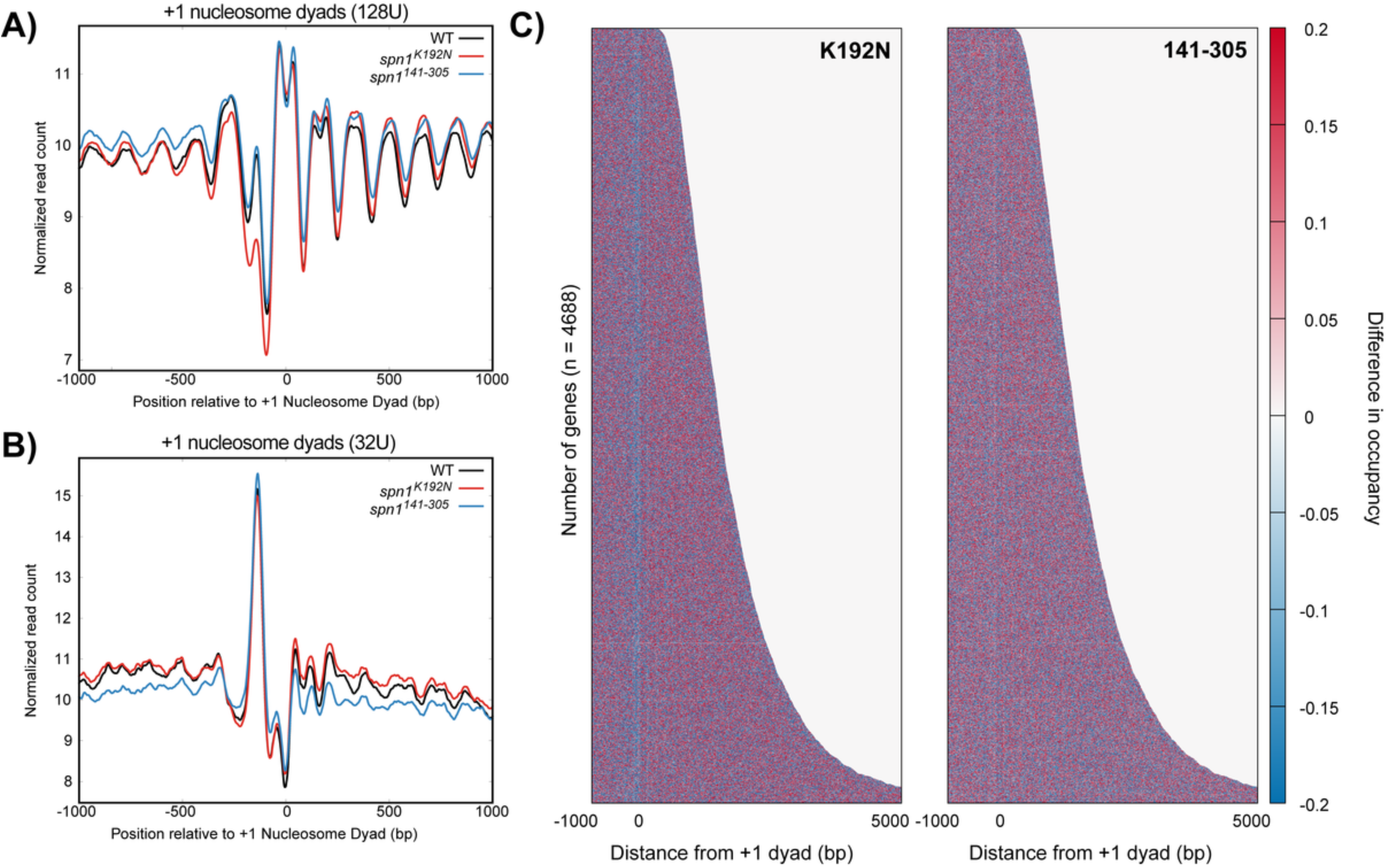
The *spn1^K192N^* mutation reduces the stability of nucleosome-depleted regions. (A) Profiles of MNase protected fragments (0-60 bp) mapped over 4688 genes aligned to the +1 nucleosome dyad in wild-type, *spn1^K192N^*, and *spn1^141-305^* expressing yeast. (B) Profiles of MNase protected fragments (0-60 bp) from low MNase-digested samples (32U) mapped over 4688 genes aligned to the +1 nucleosome dyad in wild-type, *spn1^K192N^*, and *spn1^141-305^* expressing yeast. (C) Difference heatmap between wild-type and mutant Spn1 expressing yeast strains scored by changes in occupancy of 0-60 bp fragments. Genes are aligned by their +1 nucleosome dyad position and ranked by length. Scores are calculated by subtracting wild-type occupancy data from mutant occupancy. Regions in red are enriched for fragments at that position while blue regions are depleted for fragments.

NDRs are maintained and protected by a wide range of proteins, including those sensitive to MNase (Chereji et al. 2017; Henikoff et al. 2011; Kubik et al. 2015). By varying MNase concentration, less stable structures can be revealed that are more sensitive to higher levels of MNase. To probe the MNase sensitivity of the NDR in Spn1 mutants, datasets generated from treatment with a lower amount of MNase were analyzed. We mapped smaller fragments from these samples and found that the NDR is largely unaffected in these samples (**Figure 2B**). This demonstrates that *spn1^K192N^* cells exhibit a defect in protein protections in the NDRs, which renders the protections significantly less stable than wild-type cells at higher MNase concentrations. Taken together, this indicates that the stability of protein protections in the NDR is compromised in *spn1^K192N^*cells.

The loss of short protections in the NDR prompted the question of whether longer protections are also affected in *spn1^K192N^*. To assess which lengths of protected fragments are reduced in this region, bins of fragment lengths were generated to selectively analyze their enrichment in wild-type versus Spn1 mutant-expressing yeast. Fragments approximately 147bp in length are captured in a bin of fragments between 142-152 bp in length, and can be interpreted as nucleosomal protections, as the DNA is protected from MNase by its histone contacts. These fragments represent a significant portion of the sample, as the MNase-SSP libraries exhibit a local peak in fragments of this length (**Figure S1B, File S1**). The abundance of protected fragments both 83-93 bp and 102-112 bp can be interpreted as subnucleosomal protections with loss of histone-DNA contacts due to removal of an H2A-H2B dimer (Ramachandran et al. 2017; Rhee et al. 2014). Additionally, mapping fragments of 61-71 bp can be interpreted as subnucleosomal protections with loss of histone-DNA contacts for both H2A-H2B dimers (Ramachandran et al. 2017). All of these ranges in fragment lengths also correspond to peaks in the fragment length distribution of our samples (**Figure S1B, File S1**). To determine if these protections over NDRs change upon Spn1 mutation, fragments from each of these bins were mapped over the yeast genome. The loss of fragments observed in *spn1^K192N^*diminishes as fragment length increases (**Figure S3, File S1**). This suggests that Spn1 maintains mainly short non-nucleosomal protections in the NDR.

The prior metagene analyses demonstrated that *spn1^K192N^*reduces protections over the NDR, with this defect becoming increasingly dramatic as the DNA fragment lengths shorten. To investigate the fraction of NDRs driving this effect in the metaplots, the normalized counts of 0-60 bp fragments were plotted as a heatmap, with genes aligned by their +1 nucleosome dyad position and ranked by length (**Figure S4, File S1**). To visualize the difference in fragment occupancy between wild-type and mutant strains, the wild-type occupancy scores were subtracted from the *spn1^K192N^* or *spn1^141-305^* samples to generate a difference heatmap (**Figure 2C**). In *spn1^K192N^* cells but not *spn1^141-305^* cells, reduced 0-60 bp fragment occupancy was observed directly upstream of the +1 dyad locations across genes ranked by length. This indicates that the defect in short protections in NDRs is consistent across all yeast genes in *spn1^K192N^* cells.

While nucleosomes are generally depleted in NDRs, recent work has elucidated that a subset of yeast NDR regions do contain fragile nucleosomes which are highly sensitive to MNase (Chereji et al. 2017; Kubik et al. 2015). Our analyses above demonstrate that Spn1 maintains primarily short non-nucleosomal protections in NDRs across the yeast genome. However, it is possible that the subset of NDRs containing fragile nucleosomes may exhibit larger defects in these non-nucleosomal protections compared to NDRs without fragile nucleosomes in *spn1^K192N^* cells. To address this possibility, we mapped the enrichment of short (<60 bp) fragments relative to the +1 nucleosome position of genes with either a fragile –1 nucleosome or stable –1 nucleosome as defined by Kubik *et al*. (Kubik et al. 2015). To assess the MNase sensitivity of these protections, we mapped fragments from both conditions treated with a higher or lower amount of MNase as previously described. This analysis demonstrates that both NDRs with a fragile –1 nucleosome and those with a stable –1 nucleosome experience a similar loss of 0-60 bp protections in *spn1^K192N^* cells but display no differences in nucleosomal protections from wild-type in the NDR (**Figure S5, File S1**). Taken together, this indicates that Spn1 maintenance of short non-nucleosomal protections in NDRs is critical at both of these classes of NDRs found in the yeast genome.

### Spn1 influences nucleosome positioning over genes

Spn1 is a transcription elongation factor physically associated with the RNAPII elongation complex (Ehara et al. 2022), and its genome-wide occupancy is highest over gene bodies (Mayer et al. 2010; Rossi et al. 2021). To further investigate the role of Spn1 on preserving chromatin structure over gene bodies, the normalized counts of 80-170 bp fragments were first plotted as a heatmap (**Figure S6A, File S1**). To visualize differences in pattern between wild-type and mutant strains, the wild-type occupancy scores were subtracted from the *spn1^K192N^*or *spn1^141-305^* samples to generate a difference heatmap (**Figure 3A**). As this range of fragments is sufficiently wide to include subnucleosomal protections, this analysis also shows the loss of protection across NDRs in the *spn1^K192N^* mutant strain (**Figure 3A**, left panel). The difference heatmap also reveals a striated pattern across the open reading frames of genes, which is strikingly similar in both mutants. This striated pattern of blue and red stripes seen in the difference heatmap highlight regions of lost fragment occupancy over wild-type dyad positions followed by an increase in fragment occupancy directly downstream of these positions. To specifically assess changes in nucleosome protections in the Spn1 mutant expressing strains, we repeated our analysis by exclusively mapping 142-152 bp fragments over our gene list (**Figure S6B, File S1**). We observe a striated pattern in the difference heatmap over open reading frames similar to the pattern observed in the difference heatmap generated with 80-170 bp fragments. Consistent with our prior profile analysis (**Figure S3, File S1**), we observe no loss of protection in 142-152 bp fragments over the NDR in the *spn1^K192N^*strain (**Figure S6C, File S1**).

**Figure 3.**
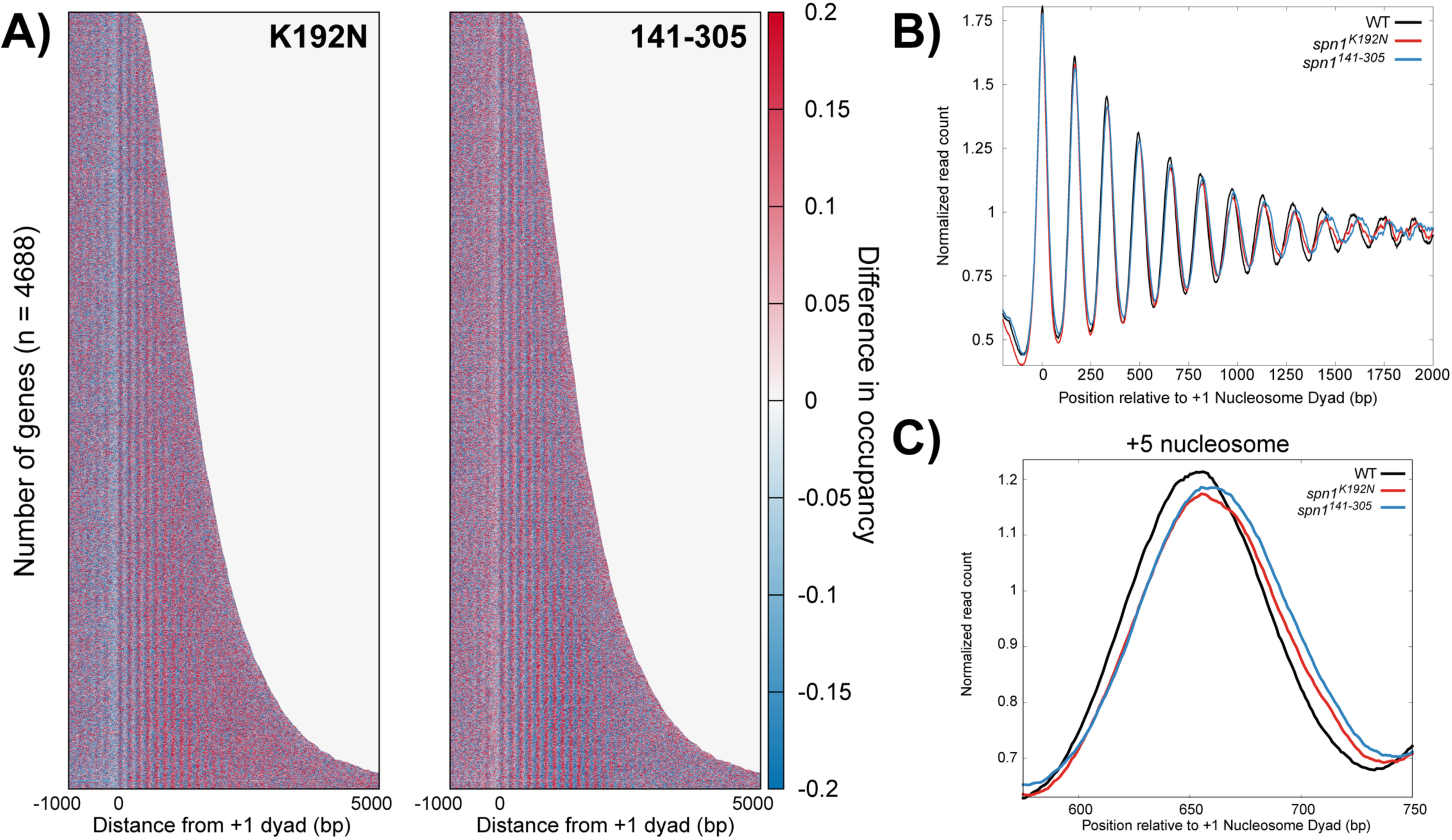
Mutations in Spn1 produce a downstream shift in nucleosome positions over open reading frames genome wide. (A) Difference heatmap between wild-type and mutant Spn1 expressing yeast strains scored by changes in nucleosome occupancy. Genes are aligned by their +1 nucleosome dyad position and ranked by length. Scores are calculated by subtracting wild-type nucleosome occupancy data from mutant nucleosome occupancy. Regions in red are enriched for nucleosomes at that position while blue regions are depleted for nucleosomes. (B) Profiles of MNase protected fragments (80-170 bp) mapped over 4688 genes aligned to the +1 nucleosome dyad in wild-type, *spn1^K192N^*, and *spn1^141-305^* expressing yeast. (C) Profiles of MNase protected fragments (80-170 bp) aligned to the +1 nucleosome dyad in wild-type, *spn1^K192N^*, and *spn1^141-305^* expressing yeast highlighting the +5 nucleosome position.

To investigate nucleosome positioning across gene bodies further, an additional metagene analysis was performed by averaging the normalized read count over genes and plotting the result as a profile. The wild-type strain produces the expected profile of regularly phased and well-positioned peaks, interpreted as peaks in nucleosome occupancy (**Figure 3B**). Nucleosome positions become “fuzzier” with each downstream nucleosome from the TSS; increases in the variability of distance of downstream nucleosome positions from the TSS result in a wider and flatter peak within the population. To quantify the differences between the wild-type and mutant profiles, a difference plot between wild-type and mutant enrichment was generated (**Figure S6D, File S1**). This difference plot indicates that nucleosome positions +1 and +2 shift specifically in cells expressing the *spn1^141-305^* allele, and further downstream nucleosome positions shift in both mutant strains. This result suggests that Spn1 maintains nucleosome positioning across genes, but the contributions of its functional domains differ in the maintenance of nucleosomes at the 5’ ends of genes versus across gene bodies.

The profiles of wild-type and mutant normalized read count indicate that that expression of either *spn1^K192N^* or *spn1^141-305^* mutant alleles produces a progressive 3’ downstream shift in nucleosome positioning. Rescaling this profile into individual profiles at different distances from the +1 dyad to feature the +1 to +6 nucleosome positions illustrates that expression of either mutant produces the trend of a progressive 3’ downstream shift in nucleosome positioning (**Figure 3C; Figure S7, File S1**). These findings align with recently published work that employed conventional MNase-seq to demonstrate Spn1 maintains nucleosome positioning over gene bodies, as expression of *spn1^K192N^* produced a progressive downstream shift in nucleosome positioning (Mayer et al. 2010; Rossi et al. 2021). The findings from our MNase-SSP approach are similar and support this notion, as well as provide the new finding that expression of the functionally distinct mutant allele *spn1^141-305^* also exhibits a defect in nucleosome positioning.

### Quantification of nucleosome positioning shifts reveals that rotational position preference is redistributed in Spn1-mutant cells particularly at longer genes

Our analysis of the MNase-SSP data from wild-type and mutant Spn1 expressing yeast thus far has revealed global shifts in chromatin protections over open reading frames. Genes across the yeast genome vary in length, and this prompted an investigation of gene length as a parameter for nucleosome shift. To begin examining this relationship, the gene list was ranked by length into quartiles of equal gene numbers (**Figure S8A, File S1**). Mapping fragments 80-170 bp and generating a difference plot as performed previously shows that the furthest downstream nucleosomes present in the longest quartile of genes exhibit the largest shift in nucleosome position (**Figure S8B, File S1**). Strikingly, the set of shortest genes experienced, on average, minimal defects in nucleosome positioning, even at nucleosome positions that were shifted in the longer genes.

To statistically validate shifts in nucleosome positioning, we developed a method to quantify the occupancy of nucleosomes at each rotational position. These positions correspond to DNA sequences with favorable histone-DNA interactions (Satchwell et al. 1986; Segal et al. 2006). Midpoints from sequenced fragments 142-152 bp in length were first mapped to the genome over each quartile ranked by gene length. This analysis reveals that peaks in 142-152 bp fragments in the mutant strains occupy the same rotational positions as wild-type cells, but with changes in occupancy at specific alternative rotational positions. Occupancy is gained downstream of the dyad and lost upstream of the dyad location in the mutant strains, indicative of a shift in nucleosome positioning (**Figure 4A; Figure S9, File S1**). As this shift manifests as a change in occupancy at discrete positions, this can be quantified by calculating the change in occupancy at each rotational position for each nucleosome. To accomplish this, Z-scores across rotational positions spanning −50 to +50 bp surrounding each nucleosome dyad position were calculated for each mapped dyad for each gene. The Z-scores of wild-type Spn1 were subtracted from the Z-scores of Spn1 mutants for each gene to obtain changes in rotation position preference due to Spn1 mutation. The median difference in Z-scores was plotted across all genes at each rotational position of each nucleosome position was plotted. The previous analysis indicates that the shifts in nucleosome positioning correlated with gene length. To evaluate the extent to which these correlations are significant, Z-scores and p-values (using the Wilcoxon Rank Sum test) were calculated for the changes in fragment midpoint occupancy at each rotational position around each nucleosome dyad for the genes in our previously determined quartiles of gene length. This analysis demonstrates that shifts in nucleosome positioning occur most dramatically at downstream nucleosome positions in longer genes, as there were more rotational positions with significant changes in Z-scores over these genes. (**Figure 4B**). Additionally, *spn1^141-305^*mutant cells exclusively have rotational positions with statistically significant changes in nucleosome occupancy at the +1 and +2 nucleosome positions. This demonstrates that the disordered N- and C-terminal regions of Spn1 specifically play a role in the proper maintenance of nucleosomes at the 5’ end of open reading frames. While both mutant strains experience shifts in nucleosome positions, *spn1^141-305^* mutant cells exhibit more positions with significant changes in 142-152 bp protected fragment occupancy over open reading frames. These results support the model that Spn1 preserves nucleosome positioning over genes, and this function becomes increasingly critical at longer genes.

**Figure 4.**
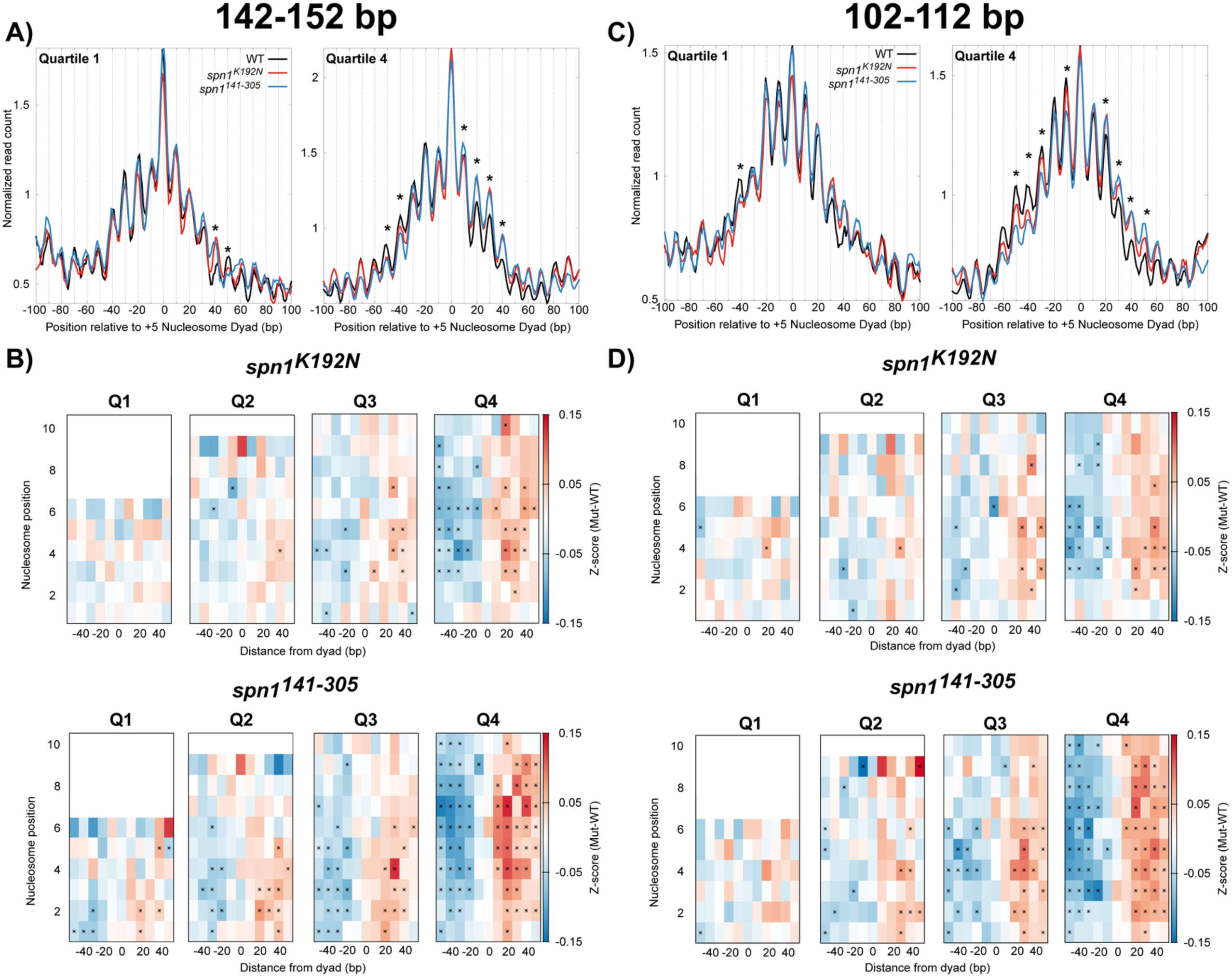
Quantification of nucleosome and subnucleosome positioning shifts in Spn1-mutant cells reveals increasing defects with increasing gene length. (A) Metagene plots of fragment midpoints 142-152 bp in length mapped to the +5 nucleosome dyad at gene quartiles ranked by length. Asterisks indicate rotational positions with statistically significant change in Z-score. (B) Z-score changes in 142-152 bp fragment occupancy at rotational positions surrounding each nucleosome dyad position at genes ranked by length in mutant Spn1 expressing yeast strains compared to wild-type plotted as a heatmap. (C) Metagene plots of fragment midpoints 102-112 bp in length mapped to the +5 nucleosome dyad at genes quartile ranked by length. Asterisks indicate rotational positions with statistically significant change in Z-score. (D) Z-score changes in 102-112 bp fragment occupancy at rotational positions surrounding each nucleosome dyad position at genes ranked by length in mutant Spn1 expressing yeast strains compared to wild-type plotted as a heatmap.

### Subnucleosomal protections are shifted in Spn1 mutant strains

An advantage to the MNase-SSP approach is the retention of small fragments in MNase-digested samples as previously described (**Figure S1B, File S1**), which can be mapped to the genome to determine the occupancy and positioning of subnucleosomal protections (Henikoff et al. 2011; Ramachandran et al. 2017). The abundance of protected fragments 102-112 bp can be interpreted as subnucleosomal protections with loss of histone-DNA contacts to an H2A-H2B dimer and the abundance of protected fragments 61-71 bp can be interpreted as subnucleosomal protections with loss of histone-DNA contacts to both H2A-H2B dimers (Ramachandran et al. 2017). To test whether Spn1 mutation leads to a shift in the location of these subnucleosomal protections, the midpoints of protected fragments 102-112 bp and 61-71 bp in length were mapped over nucleosome dyad positions at genes ranked by gene length. As observed in the V-plots, plotting the midpoints of these fragments demonstrates the existence of alternative rotational positions around mapped nucleosome dyad positions with 10 bp periodicity on either side of the dyad (**Figure 4C; Figures S10, S11, File S1**). Occupancy at these positions can be interpreted as discrete locations of subnucleosome protections with loss of contacts to an H2A-H2B dimer. In contrast to nucleosome-protected fragments, the relative occupancy of 102-112 bp and 61-71 bp protected fragments is distributed more equally across the rotational positions upstream and downstream of the dyad position. For 102-112 bp fragments, this most probably occurs since loss of contacts to a single H2A-H2B dimer shifts the location of the midpoint on the DNA protected by a hexasome (Ramachandran et al. 2017). As transcription through nucleosomes generates hexasomes with transient loss of contacts to both proximal and distal H2A-H2B dimers, the protected fragments have midpoints shifted on either side of the dyad that result in a wider distribution of hexasome-occupied rotational positions. Intriguingly, mapping 61-71 bp protected fragments revealed that rotational positions were even more equally distributed compared to the positions observed in the 102-112 bp bin of protected fragments due to the larger shifts in their midpoints.

To quantify and statistically validate the shifts observed in *spn1^K192N^* and *spn1^141-305^* cells, Z-scores and p-values were calculated at rotational positions around each mapped dyad location as described before with gene lists ranked by gene length (**Figure 4D; Figure S12, File S1**). This demonstrated that protected 102-112 bp fragments are redistributed from upstream rotational positions to downstream positions in both *spn1^K192N^* and *spn1^141-305^* cells. As seen before, more positions had significant changes in occupancy in *spn1^141-305^* cells than *spn1^K192N^*, including shifts at the +1 and +2 dyad locations seen exclusively in *spn1^141-305^* cells, with more positions experiencing significant changes in 102-112 bp fragment occupancy at longer genes (**Figure 4D**). In contrast to the 102-112 bp fragments, protections 61-71 bp in length in *spn1^K192N^* have minimal changes compared to wild-type, even at the longest genes (**Figure S12, File S1**). However, 61-71 bp protections in *spn1^141-305^* show significant shifts to downstream rotational positions as a function of gene length. Taken together, while both Spn1 mutants exhibited a length-dependent downstream shift in rotational positioning preference of subnucleosomal protections, more positions had significant changes in occupancy in *spn1^141-305^*cells than *spn1^K192N^*.

### Shifts in nucleosome and subnucleosome positioning do not exhibit a clear correlation with gene expression as assayed by RNAPII occupancy in the Spn1 mutant strains

Spn1 is part of the RNAPII elongation complex, and its occupancy over coding genes is highly correlated to RNAPII occupancy over those genes (Mayer et al. 2010; Reim et al. 2020; Viktorovskaya et al. 2021). We next assessed if a relationship exists between gene expression and the extent of nucleosome positioning defect over gene bodies in the mutant strains. Genes were ranked by Rpb3 occupancy (a subunit of RNAPII) using published NET-seq data to generate gene quartiles ranked by gene expression (Churchman and Weissman 2011). Across all quartiles, the number of genes with nucleosomes positioned at +1 through +10 was relatively consistent (**Figure S13A, File S1**), indicating gene length will not be a confounding factor in this analysis. Mapping the sequenced fragments 80-170 bp over genes ranked with the NET-seq data and generating difference plots shows a slight trend in nucleosome positioning defects between the lowest and highest quartiles, especially at the promoter-proximal nucleosome positions (**Figure S13B, File S1**). Additionally, it appears that this effect might be greater for *spn1^141-305^.*

We utilized our quantitative and statistically validated approach to determine whether nucleosome positioning defects correlate with gene expression as assayed by RNAPII occupancy. To test this possibility, the midpoints from sequenced fragments 142-152 bp were mapped to the genome over each quartile ranked by gene expression, using the quartiles ranked with NET-seq data (**Figure 5A, Figure S14, File S1**). As expected, peaks in 142-152 bp fragments in the mutant strains occupy the same rotational positions as wild-type cells, but with changes in occupancy at specific rotational positions. To quantify the shift in nucleosome positioning over gene bodies, Z-scores across rotational positions spanning −50 to +50 bp surrounding each nucleosome dyad position were calculated for each mapped dyad for each gene as described previously. Z-scores and p-values were calculated for the changes in fragment midpoint occupancy as described previously. This analysis demonstrated that shifts in nucleosome positioning do not exhibit a linear correlation with expression in the Spn1 mutant strains (**Figure 5B**). Intriguingly, for *spn1^141-305^*, the nucleosome positioning shift is more dramatic than *spn1^K192N^* with the second, third, and fourth quartiles ranked by expression are similarly impacted. Given that Spn1 is an elongation factor, we found this result surprising. For completeness, the midpoints of protected fragments 102-112 bp and 61-71 bp in length were mapped over nucleosome dyad positions at genes ranked by gene expression (**Figure 5C; Figures S15, S16, File S1**). Shifts in subnucleosomal protections were quantified with change in Z-score as described previously for genes in quartiles ranked by expression (**Figure 5D; Figure S17, File S1**). This analysis demonstrated that shifts in subnucleosome positioning do not exhibit a correlation with gene expression as assayed by RNAPII occupancy in the Spn1 mutant strains, similar to nucleosome sized fragments.

**Figure 5.**
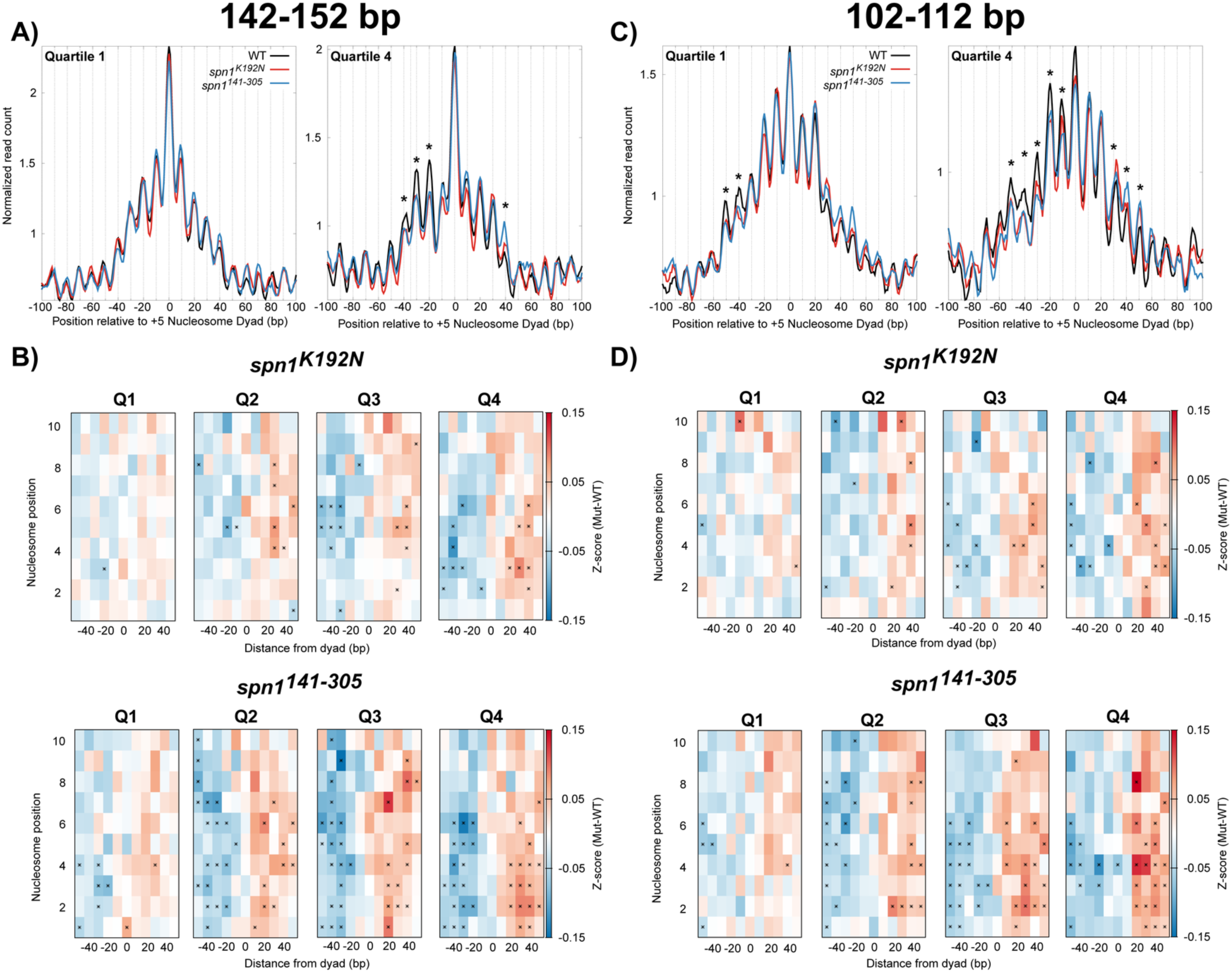
Spn1 maintains nucleosomal and subnucleosomal positioning at actively transcribed genes. (A) Metagene plots of fragment midpoints 142-152 bp in length mapped to the +5 nucleosome dyad at gene quartiles ranked by expression using published NET-seq data (38). Asterisks indicate rotational positions with statistically significant change in Z-score. (B) Z-score changes in 142-152 bp fragment occupancy at rotational positions surrounding each nucleosome dyad position at genes ranked by expression in mutant Spn1 expressing yeast strains compared to wild-type plotted as a heatmap. (C) Metagene plots of fragment midpoints 102-112 bp in length mapped to the +5 nucleosome dyad at genes quartile ranked by expression. Asterisks indicate rotational positions with statistically significant change in Z-score. (D) Z-score changes in 102-112 bp fragment occupancy at rotational positions surrounding each nucleosome dyad position at genes ranked by expression in mutant Spn1 expressing yeast strains compared to wild-type plotted as a heatmap.

### Spn1 mutant strains display global reduction in gene expression with an increased effect at longer genes

To ask if the nucleosome and subnucleosome positioning shifts correlate with changes in gene expression within the mutant strains, we performed RNA-seq on wild-type and Spn1 mutants (**File S2**). Others have shown that rapid depletion of Spn1 results in global reduction in RNA levels (Reim et al. 2020), hence we included *S. pombe* cells as a spike-in to query for the same. We performed principal component analysis of normalized transcript counts from triplicate samples of wild-type, *spn1^K192N^*, and *spn1^141-305^* expressing yeast and observed the replicates clustered together and away from replicates of the other genotypes when PC1 and PC2 loadings were plotted (**Figure 6A**). Spike-in normalization resulted in *S. pombe* genes having log_2_ fold changes (log_2_FC) distributed tightly around zero, whereas the distribution of log_2_FC of *S. cerevisiae* genes were left shifted (**Figures S18A-C, File S1**), indicating a global reduction in RNA levels. Volcano plots showed that most genes that changed significantly in *spn1^K192N^* and *spn1^141-305^* expressing yeast compared to wild-type were downregulated (**Figures 6B-D**). We combined all genes that significantly changed in at least one of the three comparisons and grouped them based on the change in patterns of expression (**Figure 6E, File S2**). Group 1 (n=821) features genes that are significantly downregulated only in *spn1^141-305^* expressing yeast compared to wild-type. Group 2 (n=280) features genes that are significantly downregulated in both *spn1^K192N^* and *spn1^141-305^* expressing yeast compared to wild-type. Group 3 (n=102) consists of genes significantly downregulated only in *spn1^K192N^* expressing yeast compared to wild-type. Group 4 (n=61) consists of genes significantly downregulated in *spn1^141-305^* expressing yeast compared to both wild-type and *spn1^K192N^* expressing yeast. The rest of the genes belong to other combinations of changes that are <20 genes per group.

**Figure 6.**
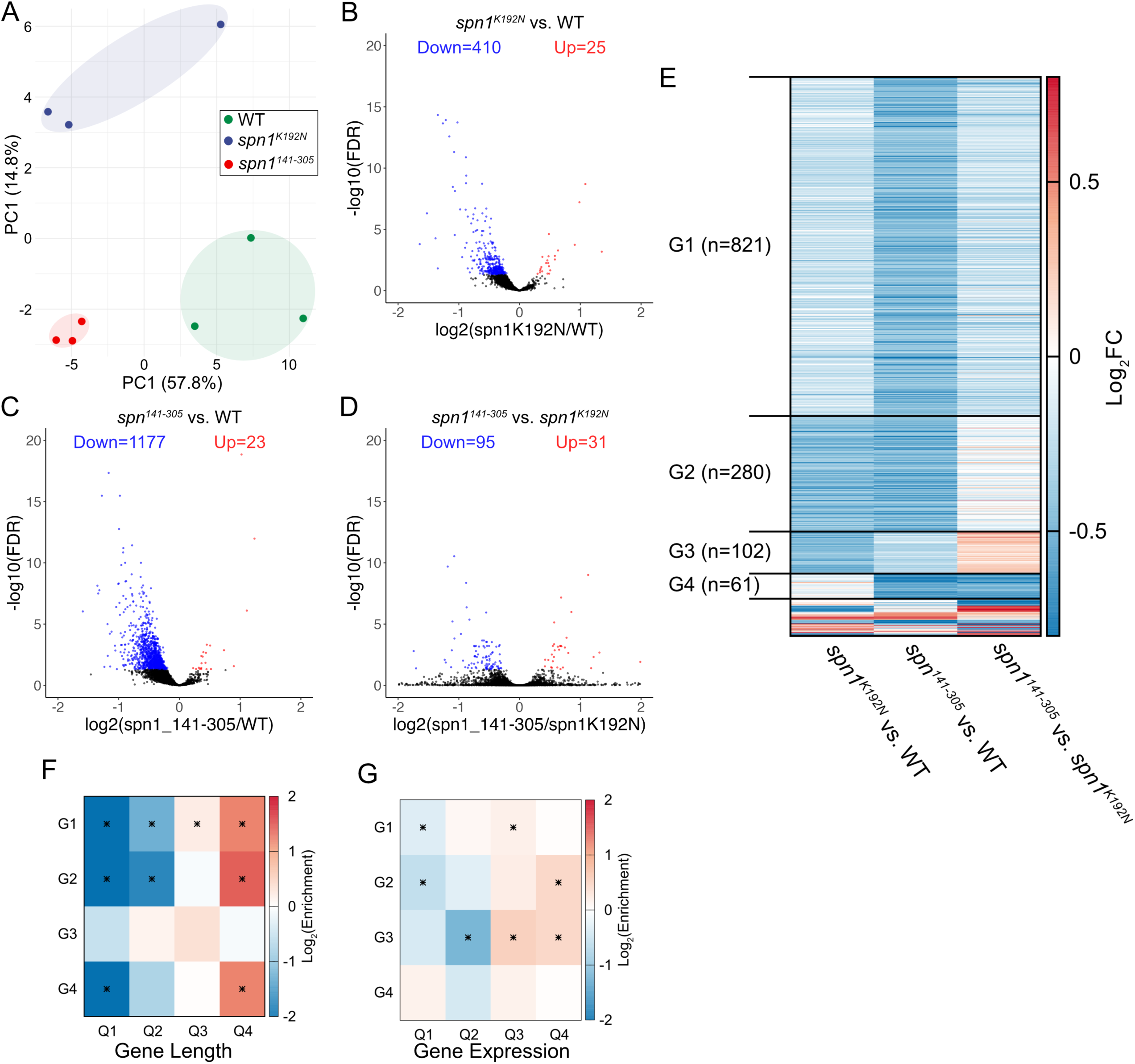
Spn1 mutants have global defects in gene expression with increased effects on longer and highly expressed genes. (**A**) Loadings of the first two principal components from the principal component analysis of normalized gene count matrix shows concordance between replicates and a clear separation between *wild-type*, *spn1^K192N^*, and *spn1^141-305^* expressing yeast. (**B**) Volcano plot where the Log_2_(Fold Change) for *spn1^K192N^* expressing yeast compared to wild-type is plotted against -log_10_(FDR). Genes with Log_2_(Fold Change) >0 and FDR ≤ 0.05 are marked in red and genes with Log_2_(Fold Change) <0 and FDR ≤ 0.05 are marked in blue. (**C**) Same as (**B**) for *spn1^141-305^* expressing yeast compared to wild-type. (**D**) Same as (**B**) for *spn1^141-305^*expressing yeast compared to *spn1^K192N^* expressing yeast. **E**) Heatmap of log_2_(Fold Change) across three possible comparisons (*spn1^K192N^* vs. wild-type*, spn1^141-305^* vs. wild-type, and *spn1^K192N^* vs. *spn1^141-305^*for all genes that have adjusted p-value<0.05 for at least one of the three comparisons. The genes are grouped (G1-G4) based on which comparisons yield significant down- or up-regulation. The rest of the genes (n=90) belong to other combinations of changes that are <20 genes per group. (**F**) Log_2_ enrichment of the overlap of gene groups defined in (D) with gene quartiles ranked by length compared to background enrichment. The statistical significance of the enrichment for each overlap was determined using hypergeometric test in R followed by multiple testing correction using the Benjamini and Hochberg method and overlaps with adjusted p-value<0.05 are marked with an asterisk. (**G**) Same as (E) for gene quartiles ranked by expression using published NET-seq data.

Our MNase-seq analysis had shown downstream nucleosome shifts that positively correlated with increasing gene length and not with genes ranked by RNAPII occupancy. We asked if the gene expression changes measured by RNA-seq have a similar correlation. Hence, we intersected lists of genes in the four groups we defined with quartiles of genes based on length and RNAPII occupancy. Strikingly, we observed a significant depletion of short genes and an enrichment of long genes in groups 1, 2, and 4 (**Figure 6E**). Intriguingly, group 3, which consists of genes that are downregulated only in *spn1^K192N^*expressing yeast compared to wild-type did not show an enrichment for long genes. Instead, group 3 genes show a clear enrichment for the highest quartile ranked by RNAPII occupancy (**Figure 6F**). Group 2 genes also show an enrichment for the highest quartile of RNAPII occupancy, indicating that *spn1^K192N^*expressing yeast have increased defects in genes with the highest gene expression quartile. We next performed gene set enrichment analysis of genes belonging to groups 1-4 and found highly significant overlap with terms related to metal ion transport, glycolysis, and ribosome biogenesis among others (**Figure S18D, File S1**).

Taken together, most genes significantly downregulated in *spn1^141-305^* or both *spn1^141-305^*and *spn1^K192N^* expressing yeast are enriched for long genes, which also correlates with the observed downstream shifts in nucleosome positioning observed for long genes. Furthermore, genes significantly downregulated in *spn1^K192N^* expressing yeast are enriched in the highest quartile of expression, which also exhibited increased downstream shifts in nucleosome positioning.

## DISCUSSION

In this study, an extensive mapping of chromatin protections in yeast expressing mutant alleles with distinct defects for known binding partners was performed to assess the role of this essential histone chaperone in maintaining chromatin structure. By retaining fragments of a wide range of sizes, the MNase-SSP approach demonstrates that Spn1 preserves nucleosomal, subnucleosomal, and non-nucleosomal protections across the yeast genome. Our analyses revealed that this function of Spn1 maintains chromatin over NDRs, the 5’ ends of genes, and within gene bodies.

The role of Spn1 in the maintenance of NDR architecture was previously unknown. While chromatin defects in Spn1 mutants have been primarily characterized over gene bodies during transcription elongation (Reim et al. 2020; Viktorovskaya et al. 2021), the *spn1^K192N^*mutation clearly reduces subnucleosomal protections in the NDR, most notably for very short protections (<60bp). How this mutation leads to this defect is unclear, although, the *spn1^K192N^*mutant is known to be defective for binding to RNAPII, likely in part through disruption of interactions with Spt5 and Spt6 (Ehara et al. 2022; Zhang et al. 2008). Also, previous work with mutants that disrupt the Spn1-Spt6 complex has implicated the complex in monitoring nucleosomes and weakening the Spn1-Spt6 interaction has impacts on the assembly and maintenance of chromatin (McCullough et al. 2015; Viktorovskaya et al. 2021). Additionally, compromise of Spt6 function has been shown to produce aberrations in chromatin over gene bodies, namely the misincorporation of histone variant H2A.Z (McCullough et al. 2015; Viktorovskaya et al. 2021). Our findings support the notion that surveillance of proper chromatin structure by the Spn1-Spt6 complex may extend to the NDRs.

Both *spn1^K192N^* and *spn1^141-305^*produced a downstream shift in nucleosome positioning over gene bodies. However, *spn1^141-305^* exclusively exhibited defects in the positioning of the +1 and +2 nucleosomes. This finding reveals that the mechanism of chromatin maintenance at the 5’ ends of genes may be functionally distinct from that over the rest of the gene and that the histone- and nucleosome-binding functions of Spn1 are part of this mechanism. As *spn1^K192N^* did not produce significant shifts in nucleosome or subnucleosome positioning at the 5’ ends of genes, one possible interpretation is that the mechanism of chromatin maintenance at the promoter-proximal region of genes requires Spn1 to directly bind chromatin, whose binding regions are unaffected in *spn1^K192N^*. In contrast, maintenance of chromatin over the rest of the body of the gene requires both Spn1 recruitment to the RNAPII complex and its histone- and nucleosome-binding capabilities, as evidenced by defects in nucleosome and subnucleosome positioning at nucleosomes distal from the promoter in both mutants.

By mapping the fragment midpoints, we utilized our dataset to demonstrate that shifts in nucleosome positioning in Spn1 mutants occur in discrete steps at alternate favored rotational positions. By quantifying the occupancy of nucleosomes at each rotational position, we present a method for statistically validating shifts in nucleosome positioning. Mapping the midpoints of smaller subnucleosomal protections also provided *in vivo* evidence suggesting the existence of alternate rotational positions occupied by subnucleosomal intermediates. These positions also occurred in discrete steps of 10 bp on either side of the dyad, and occupancy of these shorter protected fragments were distributed more evenly across the positions surrounding the mapped nucleosome dyad position. This approach of quantifying rotational positioning may be useful for other datasets of sufficiently high sequencing depth in organisms with small genomes like *S. cerevisiae*.

Quantification and statistical validation of change in preference of rotational positions at each nucleosome position of each gene allowed us to assess the relationship between the shifts in nucleosomal and subnucleosomal protections in Spn1 mutant expressing yeast and genes ranked by expression or length. As Spn1 is part of the RNAPII elongation complex, we assessed whether defects in chromatin structure observed in Spn1 were more dramatic at highly expressed genes as ranked by RNAPII occupancy. Surprisingly, our findings indicate that the relationship between transcription and changes in chromatin structure induced by Spn1 mutation were not correlated. However, Spn1 maintenance of chromatin structure became increasingly critical with gene length, as demonstrated by an increase in rotational positions with statistically significant changes in occupancy from the lowest quartile to the highest quartile of genes ranked by length. This finding provides novel insights into how chromatin structure is maintained at genes with varied length. In Spn1 mutant expressing yeast, the longest genes exhibited significant shifts in nucleosomal and subnucleosomal protections even at the most distal nucleosome positions assessed. For shorter genes, there were few significant changes in nucleosomal and subnucleosomal rotational position occupancies, even at nucleosome positions with significant changes in the longer quartiles of genes, such as the +3 and +4 positions.

To ascertain whether changes in chromatin overlap with the set of genes differentially expressed in the Spn1 mutant strains, we performed RNA-seq on wild-type and Spn1 mutant strains. This experiment revealed that yeast expressing *spn1^K192N^* or *spn1^141-305^* produce a global reduction in gene expression, with over 400 genes significantly downregulated in *spn1^K192N^*and nearly 1200 genes significantly downregulated in *spn1^141-305^*. Remarkably, we observed that these differentially expressed genes were highly enriched for long genes, the same class of genes which exhibited the largest shifts in nucleosome positioning. Additionally, we observed that genes downregulated in *spn1^K192N^* expressing yeast are enriched in highly expressed genes, which also exhibited increased shifts in nucleosome positioning.

The requirement for Spn1 in preserving nucleosome positioning shares features with the functional requirements for Spt4 and Spt6 in maintaining chromatin structure (Uzun et al. 2021; Viktorovskaya et al. 2021). Each of these factors are physically associated with RNAPII during transcription elongation, with Spn1 and Spt4 positioned at the tip of the RNAPII clamp facing the downstream DNA (Uzun et al. 2021; Viktorovskaya et al. 2021). In contrast, the main body of Spt6 is docked near the upstream DNA exit channel, although, the Spt6 N-terminal helices that bind Spn1 are proximal to the downstream DNA (Ehara et al. 2022). Spt6 binds histones and nucleosomes when the HMG-box-like protein Nhp6 is present (Ehara et al. 2022). Spt6 is commonly classified as a histone chaperone necessary for nucleosome reassembly in the wake of RNAPII, supported by its proximity to upstream DNA and its ascribed ability to suppress intragenic transcription (Doris et al. 2018; Kaplan et al. 2003; Van Bakel et al. 2013). While Spt4 has not been shown to bind chromatin, it oscillates on and off RNAPII along with its binding partner Spt5 based on nucleosome positions, and its deletion produces a downstream nucleosome positioning shift. Moreover, Spt5 deletion produces increased stalling and backtracking during early transcription elongation (Uzun et al. 2021). Despite differences in their transcription-associated functions, Spn1, Spt4, and Spt5 are all critical for maintaining proper chromatin structure and their relationships are further supported in that defects in each produced a similar pattern of downstream shifts in nucleosome positioning. One explanation for this commonality is that all participate in this process of co-transcriptional nucleosome disassembly and reassembly, and a defect in any step of this process produces a similar defect in chromatin. Others have posed that this defect in passage through nucleosomes may produce an elongation complex that transcribes through chromatin more slowly than in wild-type cells, which might produce mispositioned nucleosomes (Uzun et al. 2021). Indeed, there is evidence that loss of RNAPII transcription results in shifts in nucleosome positions, as depletion of Rpb1 (Tramantano et al. 2016) and growth of the temperature-sensitive *rpb1-1* allele at a non-permissive temperature (Tramantano et al. 2016), produces a downstream nucleosome shift.

Maintenance of nucleosome positioning is also associated with ATP-dependent nucleosome remodelers, which position nucleosomes and evict or exchange histones in an ATP-dependent manner (reviewed in (Clapier et al. 2017)). Recent work demonstrated that the ISW1a nucleosome remodeler complex subunit *Ioc4* mutant or depletion produces a downstream shift in nucleosome positioning comparable to the shift we observed in the Spn1 mutants (Bhardwaj et al. 2020). However, other MNase-seq experiments demonstrate that depletion or deletion of nucleosome remodeling factors, notably Isw1, Chd1, Ino80, and Sth1 produce upstream shifts in nucleosome positioning over gene bodies (Klein-Brill et al. 2019; Ocampo et al. 2016). Spn1 does collaborate with the SWI/SNF nucleosome remodeler complex, as we’ve previously observed through a significant genetic interaction with *SNF2*, which encodes the catalytic subunit of SWI/SNF (Zhang et al. 2008).Intriguingly, degradation of Snf2 has little effect on nucleosome positioning (Klein-Brill et al. 2019). Taken together, the shifts observed in the Spn1 mutants more closely resemble the shifts produced by Spt4, Spt6, and Rpb1 rather than most nucleosome remodelers.

A puzzling question that remains is how *spn1^K192N^* and *spn1^141-305^* produce such similar changes in chromatin protections over nucleosome positions distal to promoters despite their distinct defects for known binding partners. Both Spn1 recruitment to the elongating RNAPII complex and its ability to bind histones and/or nucleosomes are critical for preserving chromatin structure, and likely occur co-transcriptionally. Wild-type Spn1 is poised near the downstream nucleosome in the RNAPII elongation complex (Ehara et al. 2022). Spn1 directly binds histones and nucleosomes, a function that may be important for the disassembly of downstream nucleosomes during transcription. If the recruitment of Spn1 to sites of transcription is diminished, the process of disassembling the downstream nucleosome may be impacted. Conversely, the *spn1^141-305^* mutant is likely still recruited to sites of transcription. However, its loss of histone- and nucleosome-binding regions renders it unable to participate in the process of downstream nucleosome disassembly. In either mutant, the loss of Spn1 function results in defects in chromatin preservation during transcription elongation, resulting in a shift in nucleosome and subnucleosome protections. Taken together, these findings deepen our understanding of Spn1-dependent chromatin maintenance over the yeast genome and provide insight into mechanisms underlying this process.

## AUTHOR CONTRIBUTIONS

**Andrew J. Tonsager**: Conceptualization, Formal analysis, Investigation, Methodology, Visualization, Writing—original draft. **Alexis Zukowski**: Formal analysis, Methodology, Investigation. **Catherine A. Radebaugh**: Conceptualization, Formal analysis, Investigation, Methodology, Visualization. **Abigail Weirich**: Investigation. **Laurie A. Stargell**: Conceptualization, Funding Acquisition, Project administration, Supervision, Writing—review & editing. **Srinivas Ramachandran**: Conceptualization, Formal analysis, Funding Acquisition, Investigation, Methodology, Project administration, Supervision, Visualization, Writing—review & editing.

## ACKNOWLEDGEMENTS

We thank Sarah Swygert for her expert technical assistance.

