## Supplementary material for "The Histone Chaperone Spn1 Preserves Chromatin Protections at Promoters and Nucleosome Positioning in Open Reading Frames": File S1

**SUPPLEMENTARY DATA**

**Supplemental Table S1:** Plasmids and yeast strains used in this study.

| Plasmid | Description | |
|---|---|---|
| pCR311 | Full length wild type *SPN1* with 403 bp of upstream sequence and 116bp of downstream sequence, myc2 tagged at the amino terminus, pRS313 (CEN, *HIS3*) | |
| pCR312 | *spn1*$^{K192N}$ with 403 bp of upstream sequence and 116bp of downstream sequence, myc2 tagged at the amino terminus, pRS313 (CEN, *HIS3*) | |
| pAA344 | *spn1*$^{141\text{-}305}$ with 403 bp of upstream sequence and 116bp of downstream sequence, myc2 tagged at the amino terminus, pRS313 (CEN, *HIS3*) | |
| **Strain** | **Genotype** | **Source** |
| BY4741 | *MATa his3Δ1 leu2Δ0 met15Δ0 ura3Δ0* | Research Genetics |
| LZ0-1 | BY4741 *spn1Δ::LEU2* + pCR311 | (Li et al. 2018), (Zhang et al. 2008) |
| LZ0-2 | BY4741 *spn1Δ::LEU2* + pCR312 | (Li et al. 2018), (Zhang et al. 2008) |
| LZ0-3 | BY4741 *spn1Δ::LEU2* + pAA344 | (Li et al. 2018), (Zhang et al. 2008) |

**SUPPLEMENTAL FIGURES**

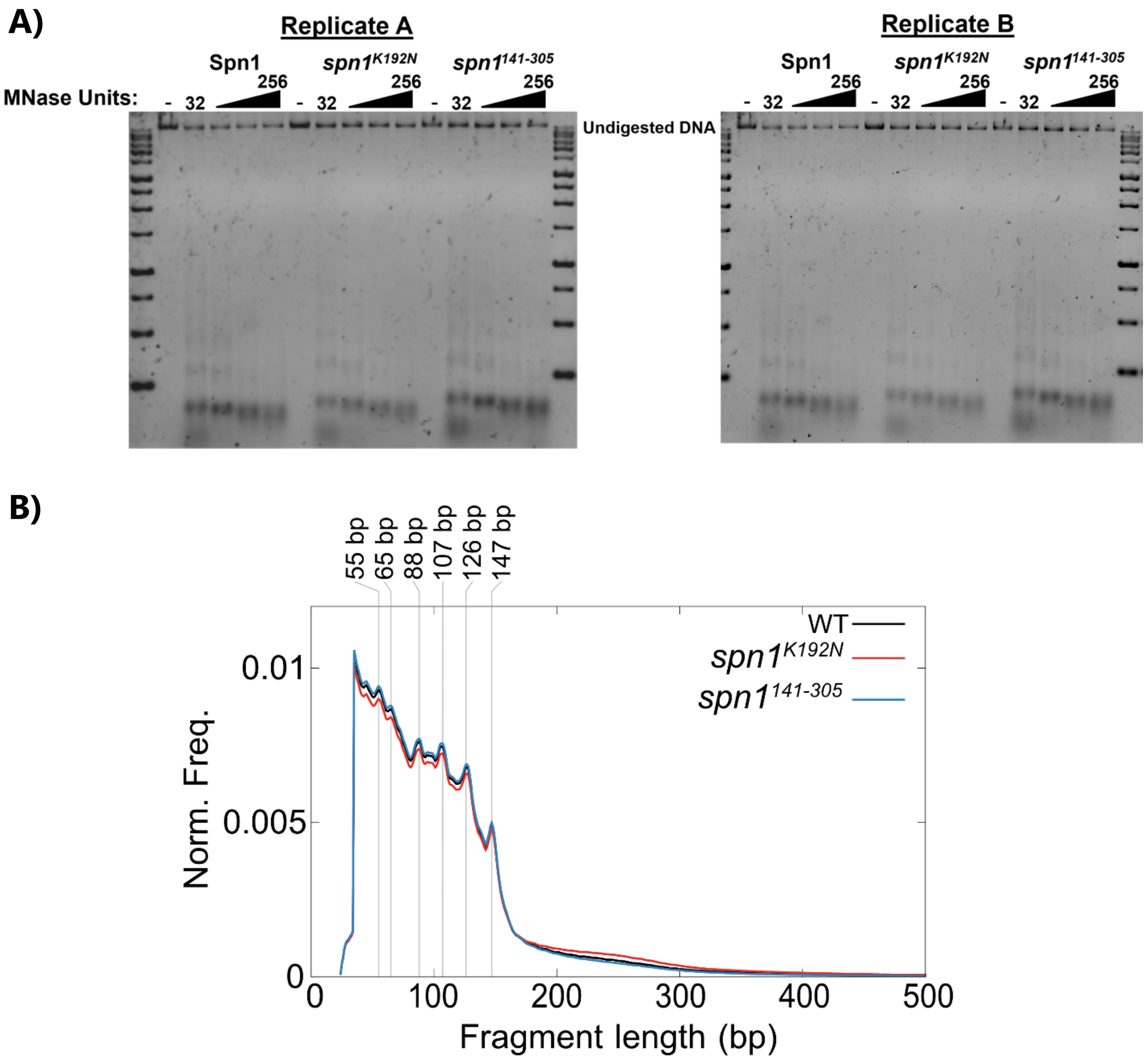


**Supplemental Figure S1: Characterization of MNase digested samples demonstrates uniform distribution of protected fragments.** (A) Yeast nuclei isolated from yeast expressing wild-type Spn1, *spn1*$^{K192N}$ and *spn1*$^{141\text{-}305}$ were digested with increasing amounts of MNase then subjected to agarose gel electrophoresis. (B) The normalized fragment length distribution of sequencing datasets generated from MNase digested samples (wild-type: black, *spn1*$^{K192N}$: red, *spn1*$^{141\text{-}305}$: blue). Fragment length distributions were normalized by averaging distributions from each sample followed by resampling.

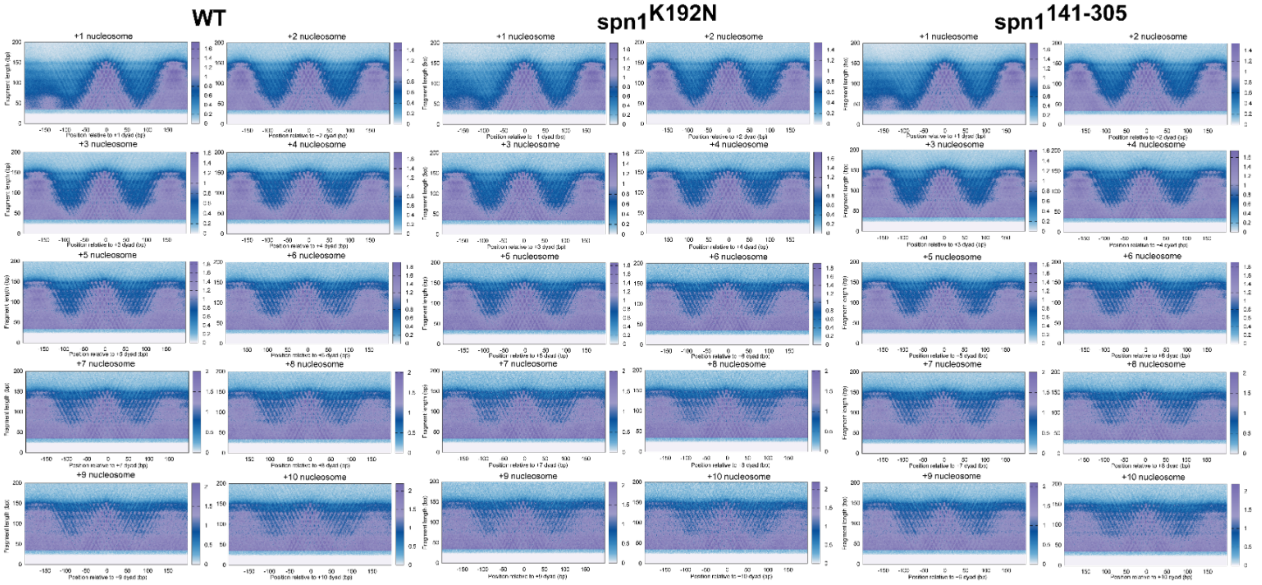


**Supplemental Figure S2:** V-plots were produced by mapping fragment midpoints at each nucleosome dyad position from positions +1 to +10 over 4688 genes in wild-type or Spn1 mutant expressing yeast.

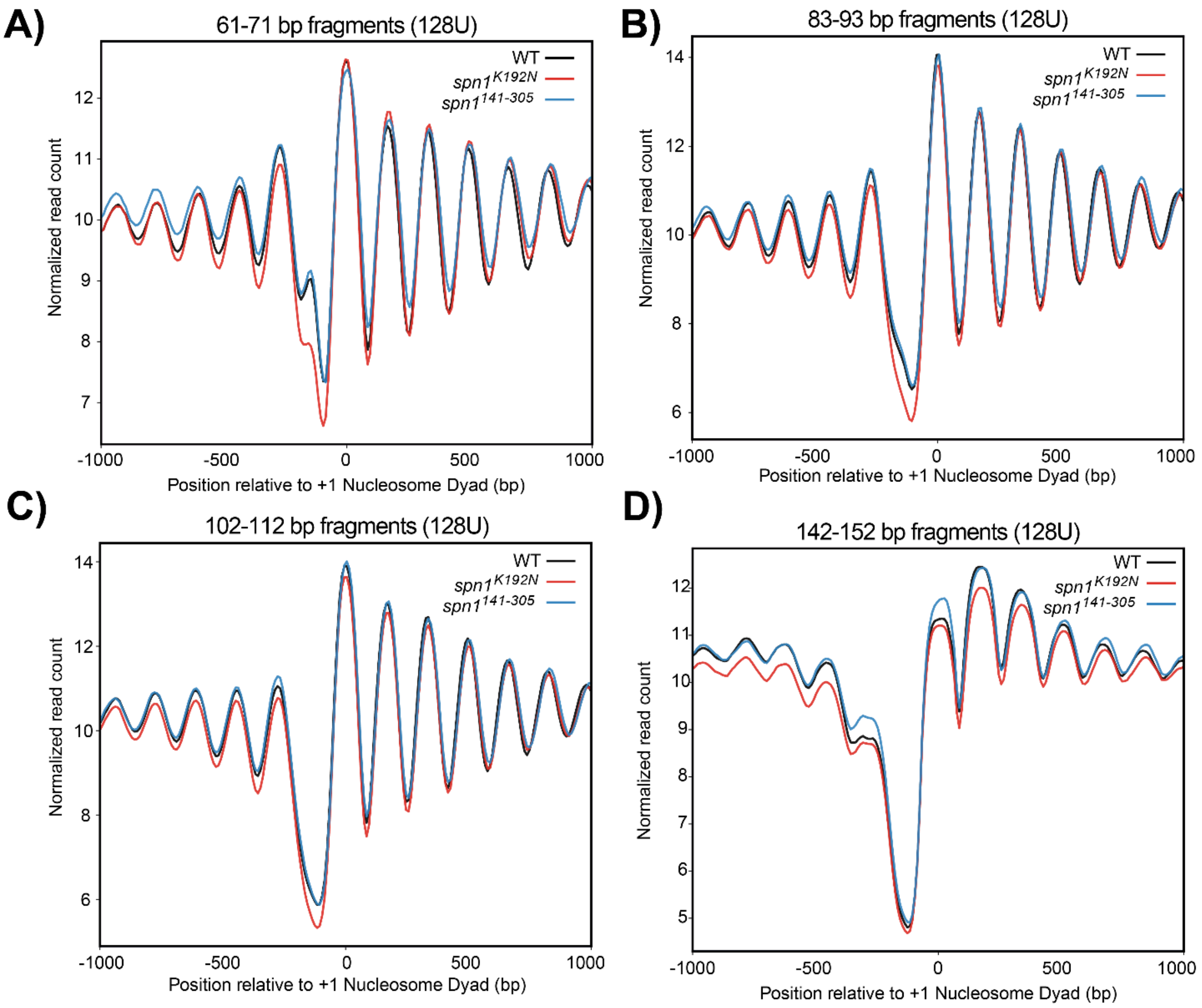


**Supplemental Figure S3: *spn1$^{K192N}$* cells exhibit a loss of MNase-protected fragments in NDRs, which diminishes as fragment length increases.** Profiles of MNase protected fragments (A) 61-71 bp, (B) 83-93 bp, (C) 102-112 bp, and (D) 142-152 bp in length mapped over 4688 genes aligned to the +1 nucleosome dyad in wild-type, *spn1$^{K192N}$*, and *spn1$^{141\text{-}305}$* expressing yeast.

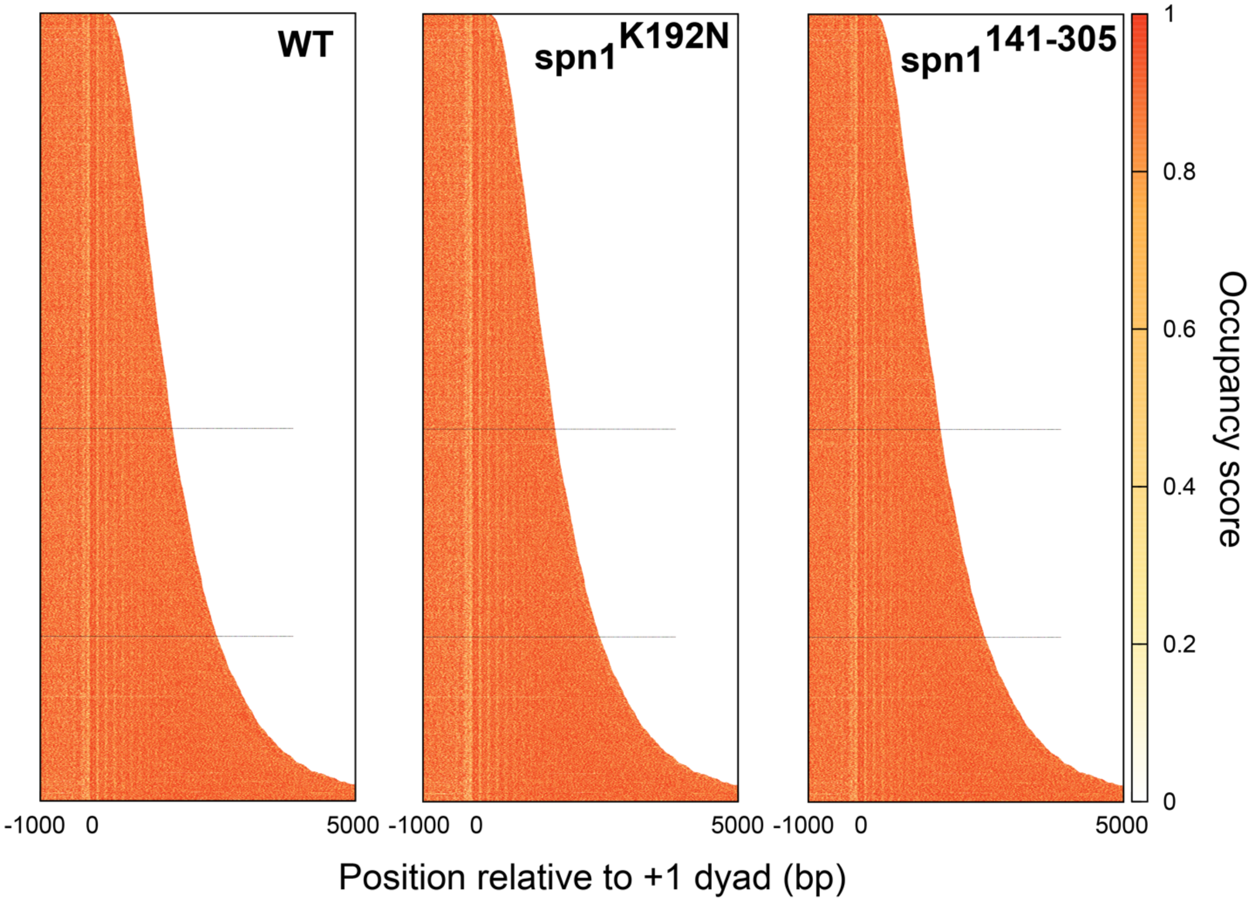


**Supplemental Figure S4: *spn1$^{K192N}$* cells experience a loss in short chromatin protections over NDRs genome wide.** Heatmaps displaying occupancy of MNase protected fragments (0-60bp) across 4688 genes in wild-type, *spn1$^{K192N}$*, and *spn1$^{141\text{-}305}$* expressing yeast.

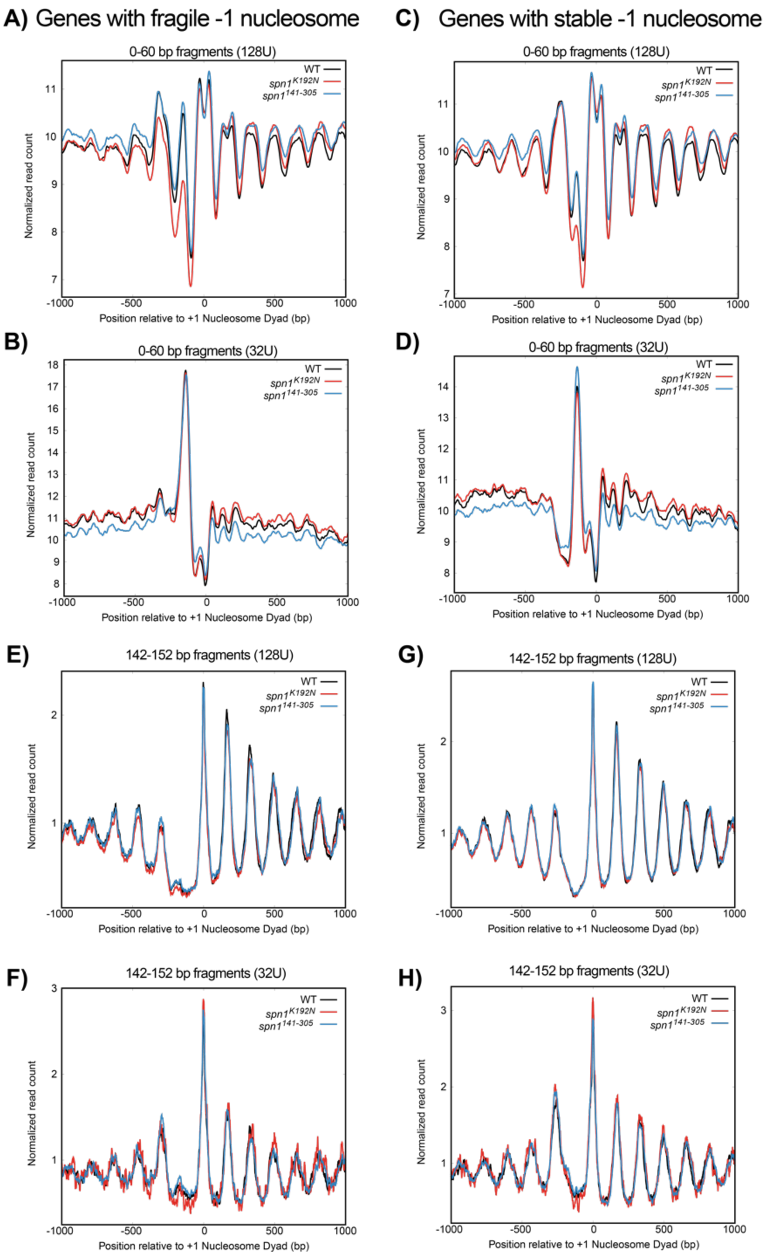


**Supplemental Figure S5:** ***spn1**$^{K192N}$* **cells exhibit a loss of MNase-protected fragments in NDRs with or without fragile nucleosomes.** Profiles of MNase protected fragments 0-60 bp in length from (A) high MNase (128U) and (B) low MNase (32U) mapped over genes containing a fragile –1 nucleosome in NDRs as defined in (Kubik et al. 2015) aligned to the +1 nucleosome dyad in wild-type, *spn1*$^{K192N}$, and *spn1*$^{141\text{-}305}$ expressing yeast. Profiles of MNase protected fragments 0-60 bp in length from (C) high MNase (128U) and (D) low MNase (32U) mapped over genes containing a stable –1 nucleosome in NDRs as defined in (Kubik et al. 2015) aligned to the +1 nucleosome dyad in wild-type, *spn1*$^{K192N}$, and *spn1*$^{141-305}$ expressing yeast. Profiles of MNase protected fragments 142-152 bp in length from (E) high MNase (128U) and (F) low MNase (32U) mapped over genes containing a fragile –1 nucleosome in NDRs aligned to the +1 nucleosome dyad in wild-type, *spn1*$^{K192N}$, and *spn1*$^{141-305}$ expressing yeast. Profiles of MNase protected fragments 142-152 bp in length from (G) high MNase (128U) and (H) low MNase (32U) mapped over genes containing a stable –1 nucleosome in NDRs aligned to the +1 nucleosome dyad in wild-type, *spn1*$^{K192N}$, and *spn1*$^{141-305}$ expressing yeast.

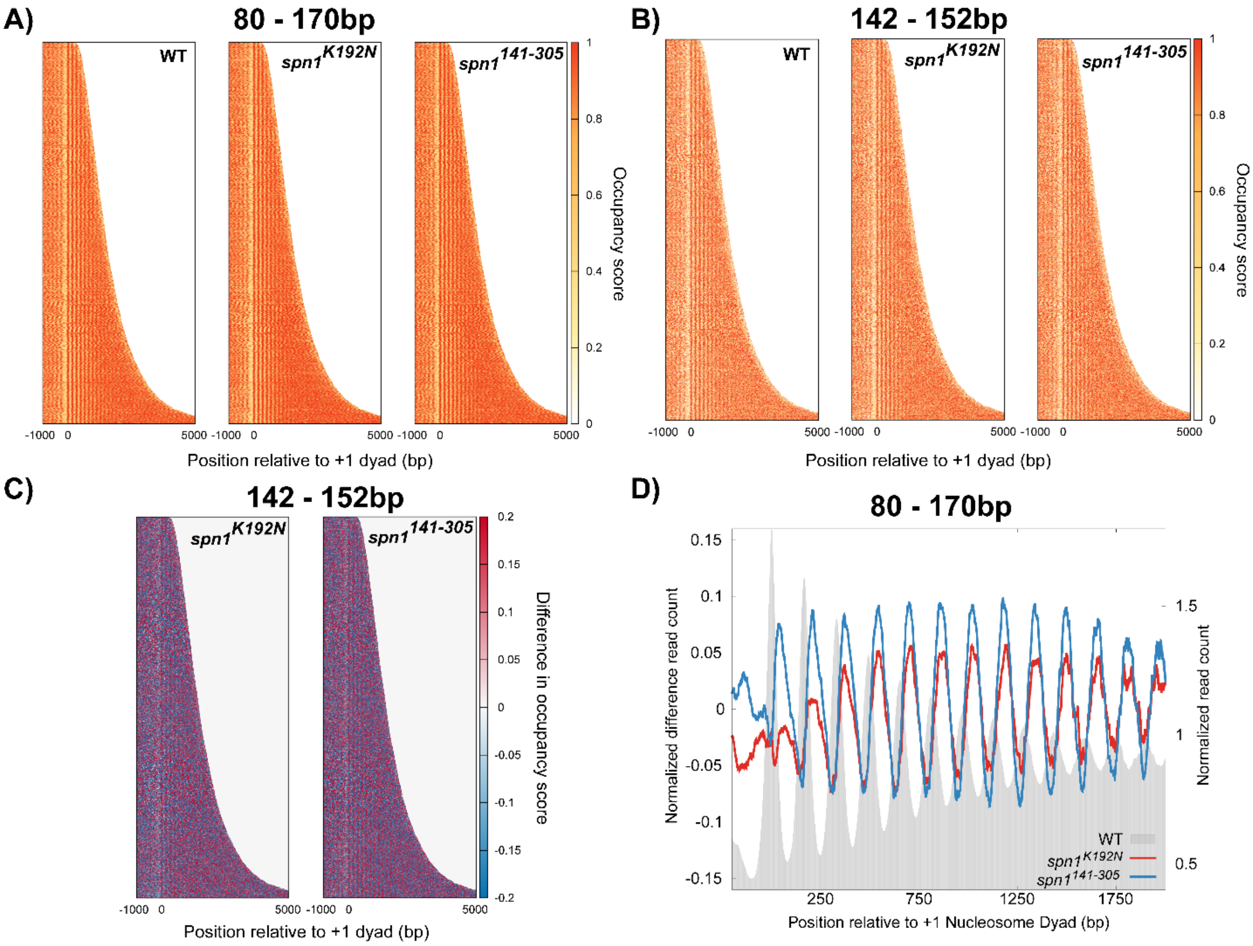


**Supplemental Figure S6: Mutations in Spn1 propagate a downstream shift in chromatin protections over open reading frames genome wide.** (A) Heatmaps of occupancy of MNase protected fragments (80-170bp) across 4688 genes in wild-type, *spn1*$^{K192N}$, and *spn1*$^{141\text{-}305}$ expressing yeast. (B) Heatmap of occupancy of MNase protected fragments (142-152bp) across 4807 genes in wild-type, spn1$^{K192N}$, and spn1$^{141\text{-}305}$ expressing yeast. (C) Difference heatmap between wild-type and mutant Spn1 expressing yeast strains scored by changes in 142-152 bp fragment occupancy. Genes are aligned by their +1 nucleosome dyad position and ranked by length. Scores are calculated by subtracting wild-type occupancy data from mutant occupancy. Regions in red are enriched for fragments at that position while blue regions are depleted for fragments. (D) Difference profile plot between wild-type and mutant samples. Average nucleosome density from wild-type expressing yeast is plotted in grey. Differences between wild-type and mutants (*spn1*$^{K192N}$, red and *spn1*$^{141\text{-}305}$, blue) against wild-type replicates are plotted.

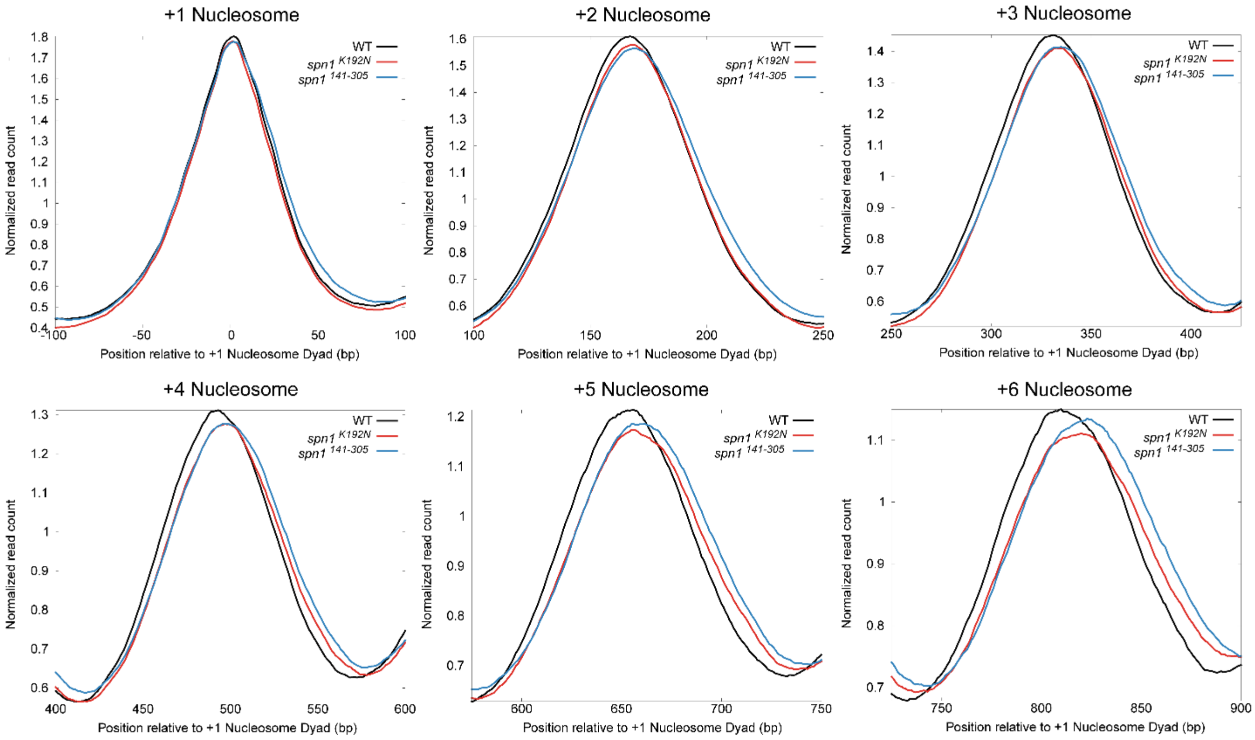


**Supplemental Figure S7: Expression of either *spn1*$^{K192N}$ and *spn1*$^{141\text{-}305}$ mutant alleles produced a progressive 3′ downstream shift in nucleosome positioning.** Profiles of MNase protected fragments (80-170 bp) mapped over 4688 genes aligned to the +1 nucleosome dyad in wild-type, *spn1*$^{K192N}$, and *spn1*$^{141\text{-}305}$ expressing yeast. Panels are rescaled by normalized read count and separated into panels to feature +1 to +6 nucleosome positions.

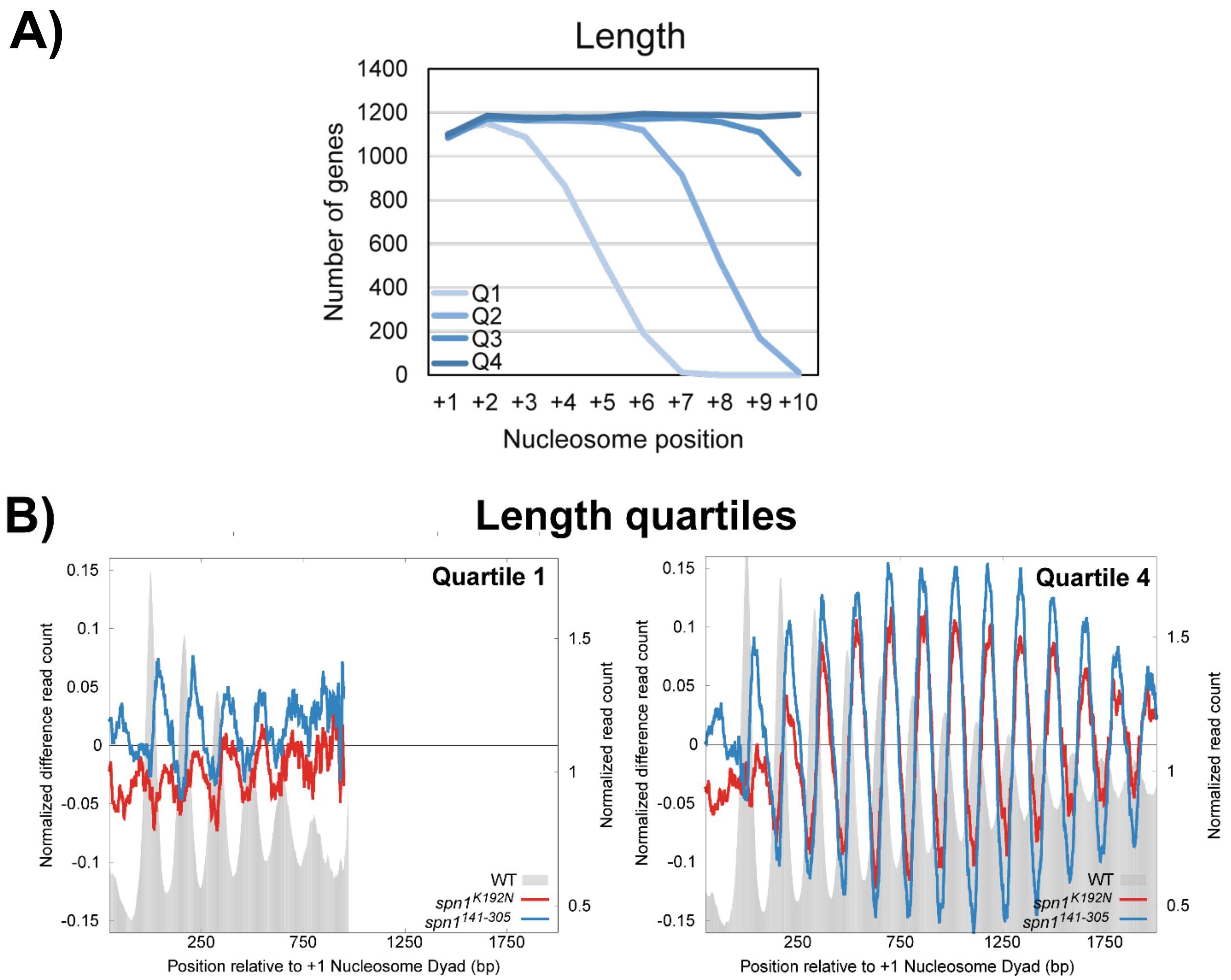


**Supplemental Figure S8: Yeast genes were ranked by gene length to generate quartile gene lists.** (A) Count of genes containing mapped nucleosomes at each position across quartiles ranked by length shows a uniform nucleosome count across quartiles at each nucleosome position. (B) Difference profile plots between wild-type and mutant samples over 4688 genes binned into quartiles by gene length, with Q1 representing the shortest genes (left panel) and Q4 representing the longest (right panel).

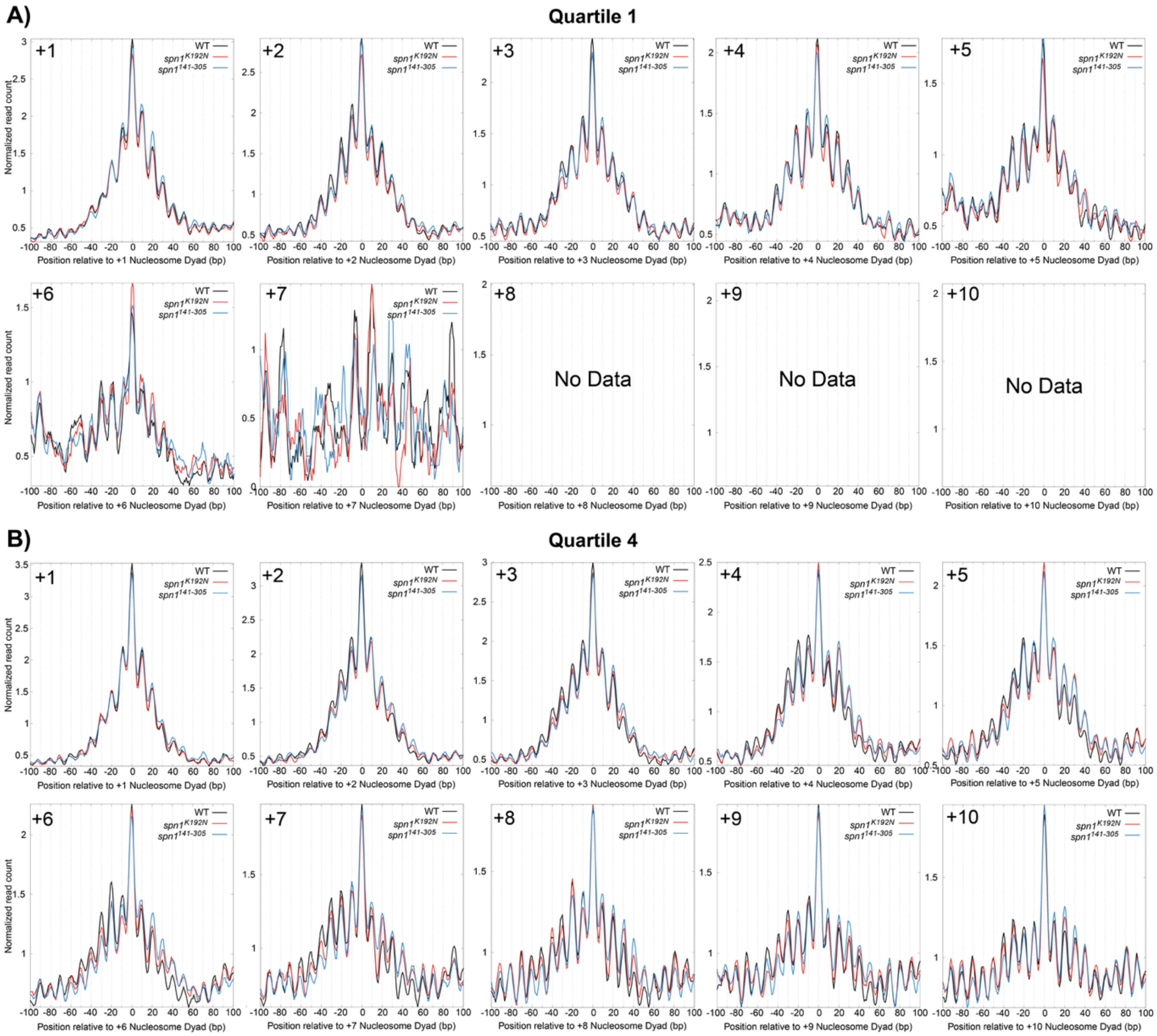


**Supplemental Figure S9: Nucleosomal protections in Spn1 mutants occupy the same rotational positions as wild-type cells, but with higher occupancy at rotational positions downstream of the dyad with a loss of occupancy at positions upstream of the dyad location.** Metagene plots of fragment midpoints 142 - 152bp in length mapped to each nucleosome dyad at the (A) shortest and (B) longest gene quartiles ranked by length.

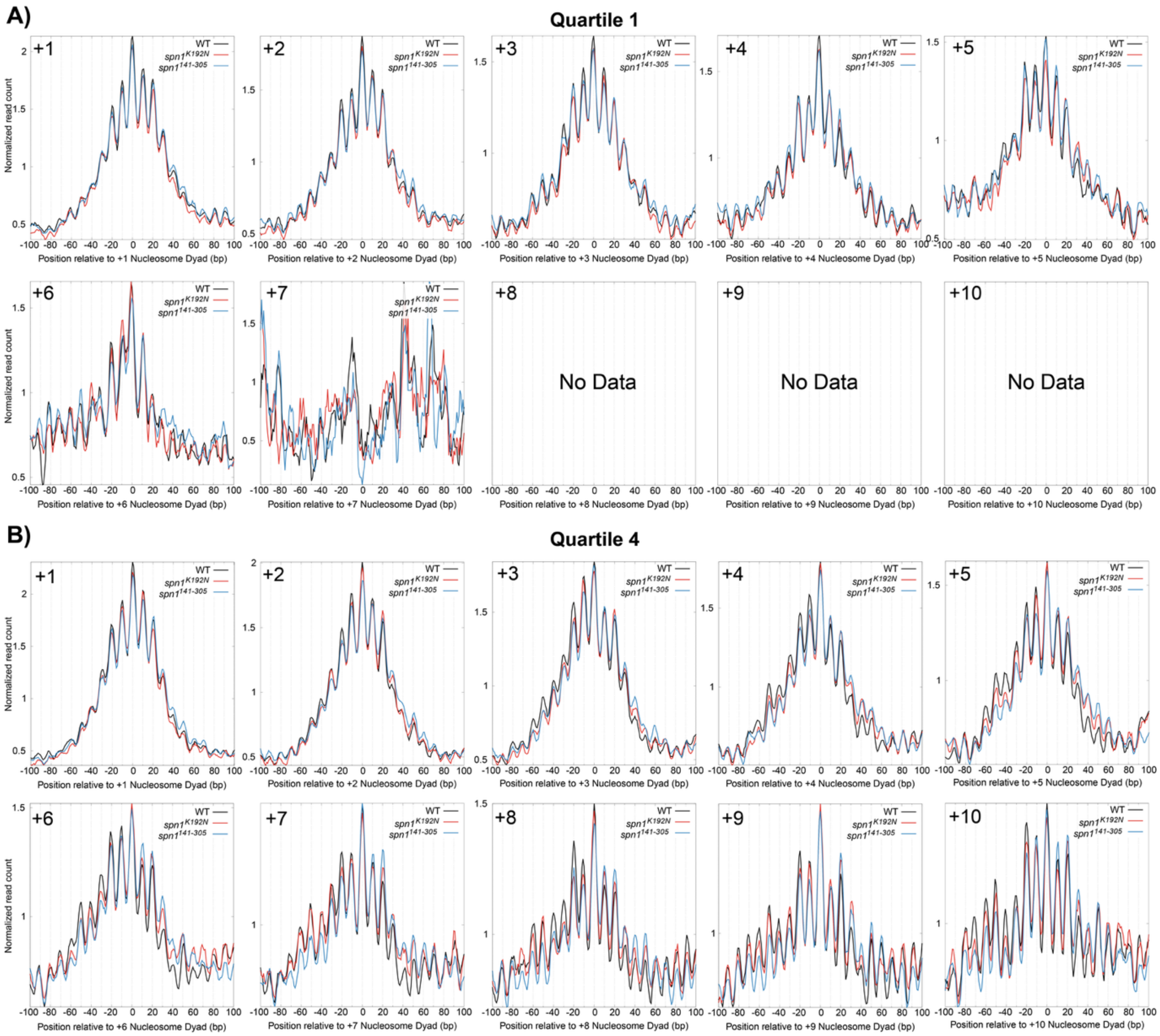


**Supplemental Figure S10: Subnucleosomal protections 102-112 bp in length in Spn1 mutants occupy the same rotational positions as wild-type cells, but with higher occupancy at rotational positions downstream of the dyad with a loss of occupancy at positions upstream of the dyad location.** Metagene plots of fragment midpoints 102 - 112bp in length mapped to each nucleosome dyad at the (A) shortest and (B) longest gene quartiles ranked by length.

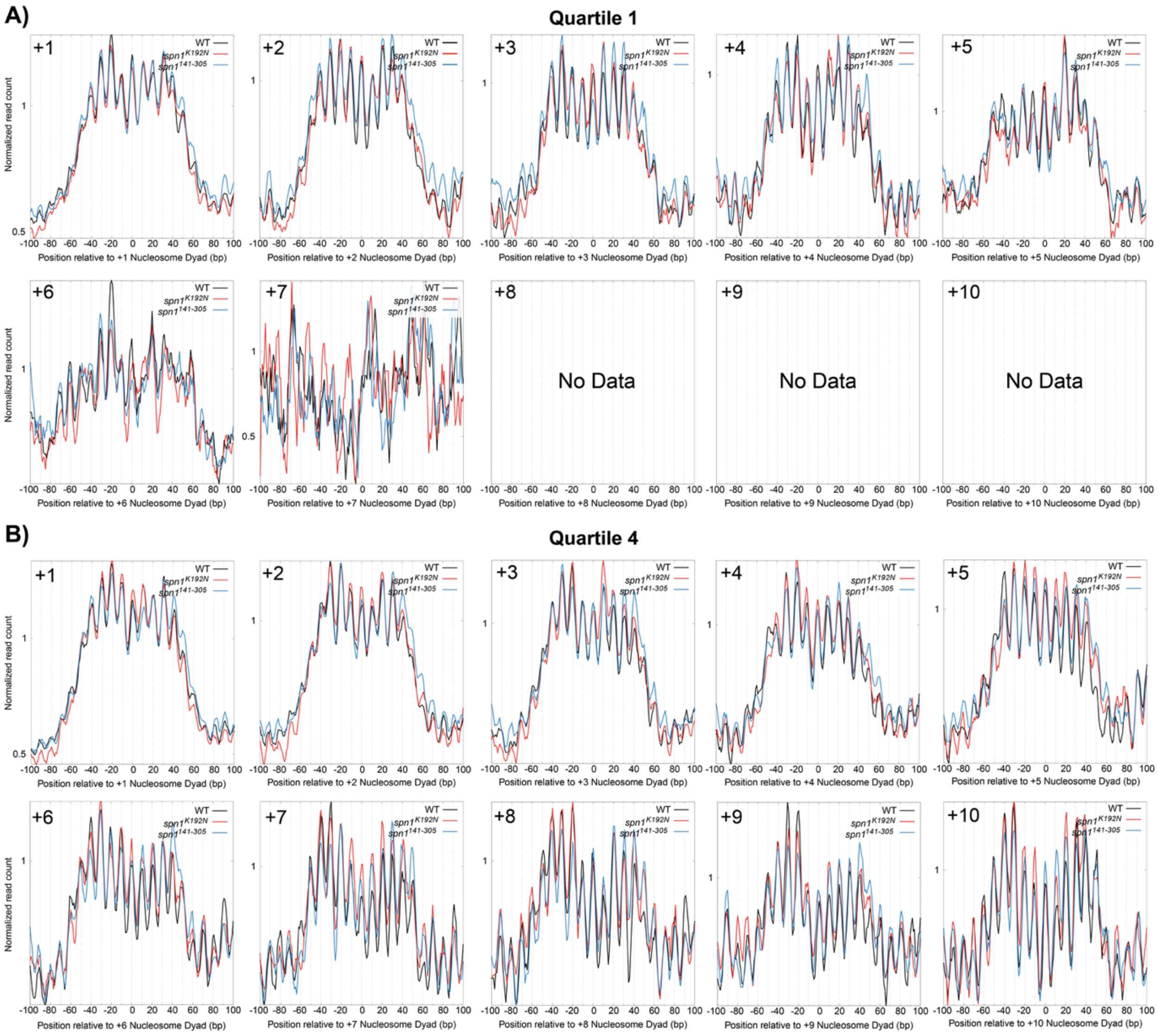


**Supplemental Figure S11: Subnucleosomal protections 61-71 bp in length in Spn1 mutants occupy the same rotational positions as wild-type cells, but with higher occupancy at rotational positions downstream of the dyad with a loss of occupancy at positions upstream of the dyad location in *spn1$^{141\text{-}305}$*.** Metagene plots of fragment midpoints 61 - 71bp in length mapped to each nucleosome dyad at the (A) shortest and (B) longest gene quartiles ranked by length.

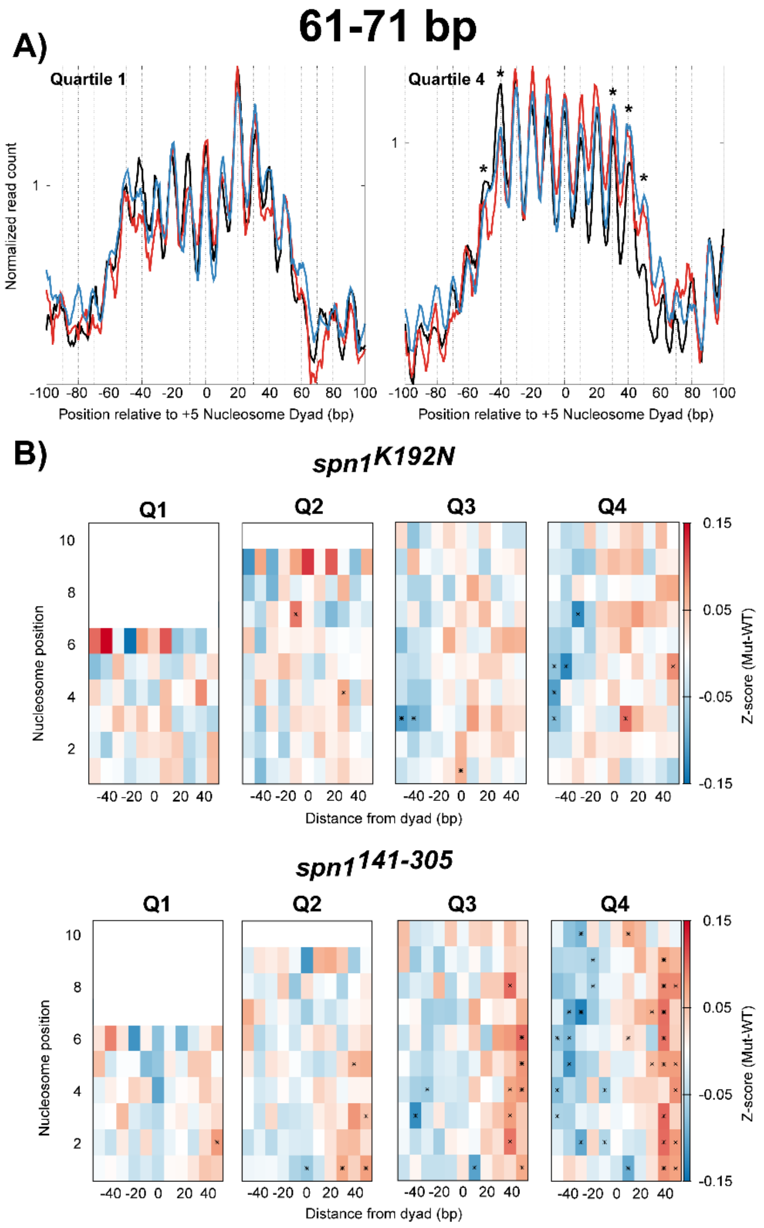


**Supplemental Figure S12: Protected fragments 61-71 bp in length experience little shifts in Spn1 mutant expressing strains.** (A) Metagene plots of fragment midpoints 61-71 bp in length mapped to the +5 nucleosome dyad at genes ranked by length. Asterisks indicate rotational positions with statistically significant change in Z-score. (B) Z-score changes in 61-71 bp fragment occupancy at rotational positions surrounding each nucleosome dyad position at genes ranked by length in mutant Spn1 expressing yeast strains compared to WT plotted as a heatmap. Asterisks indicate rotational positions with statistically significant change in Z-score.

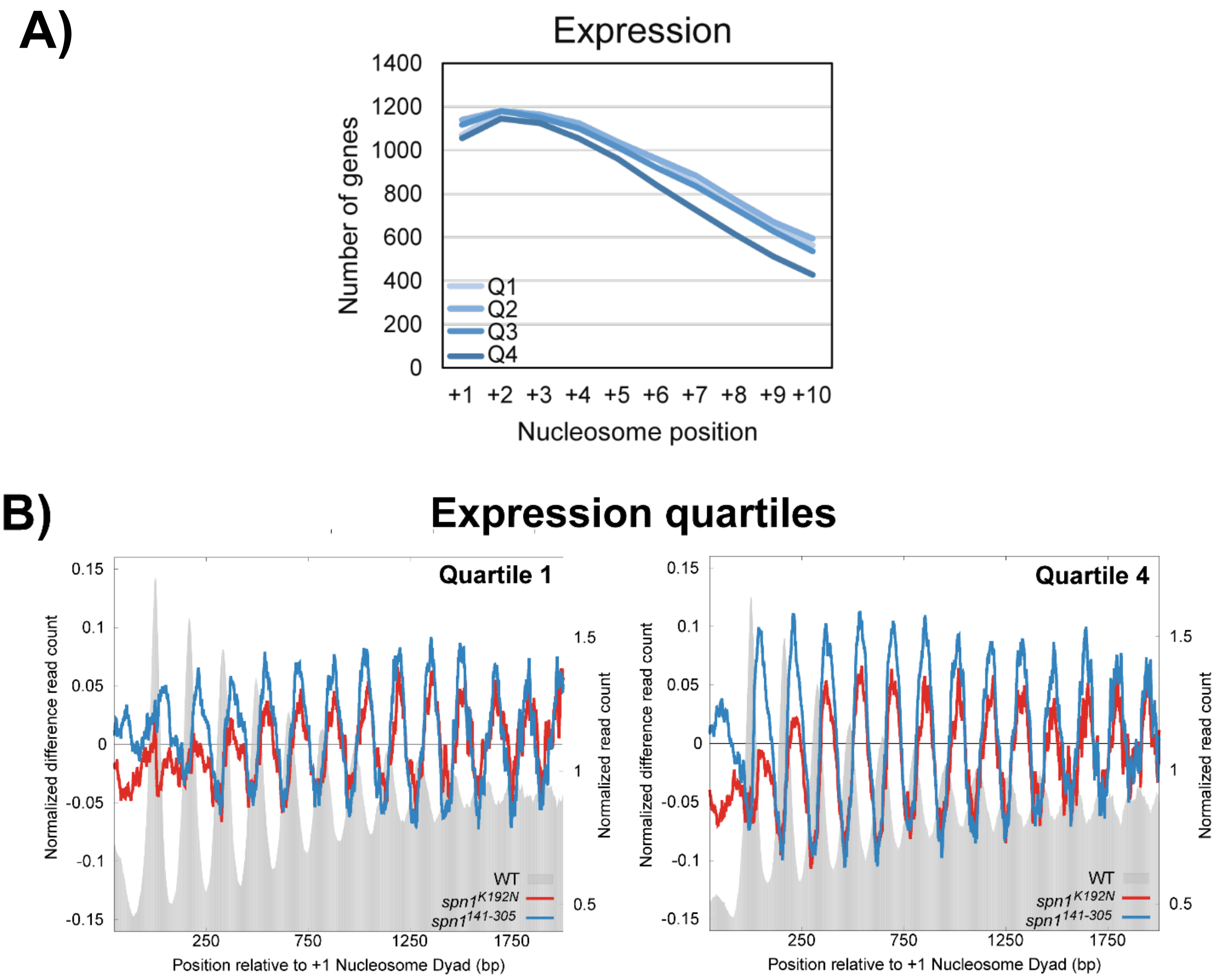


**Supplemental Figure S13: Yeast genes were ranked by published NET-seq data to generate quartile gene lists.** (A) Count of genes containing mapped nucleosomes at each position across quartiles ranked by expression shows the loss of downstream nucleosome positions in quartiles. (B) Difference profile plots between wild-type and mutant samples over 4688 genes binned into quartiles by expression, with Q1 representing the least expressed (left panel), and Q4 representing the most expressed genes (right panel).

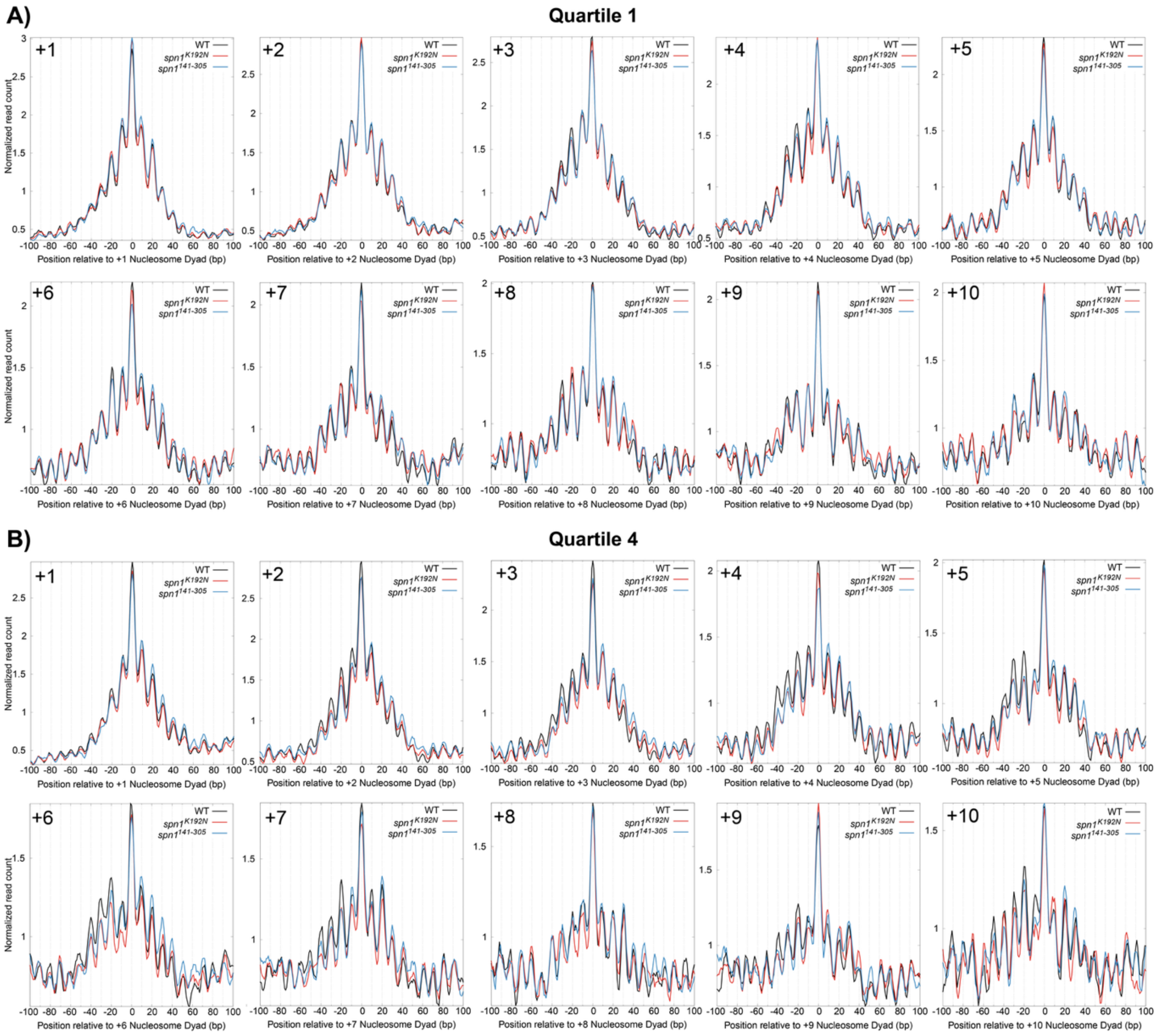


**Supplemental Figure S14: Nucleosomal protections in Spn1 mutants occupy the same rotational positions as wild-type cells, but with higher occupancy at rotational positions downstream of the dyad with a loss of occupancy at positions upstream of the dyad location.** Metagene plots of fragment midpoints 142 - 152bp in length mapped to each nucleosome dyad at the (A) lowest and (B) highest gene quartiles ranked by expression.

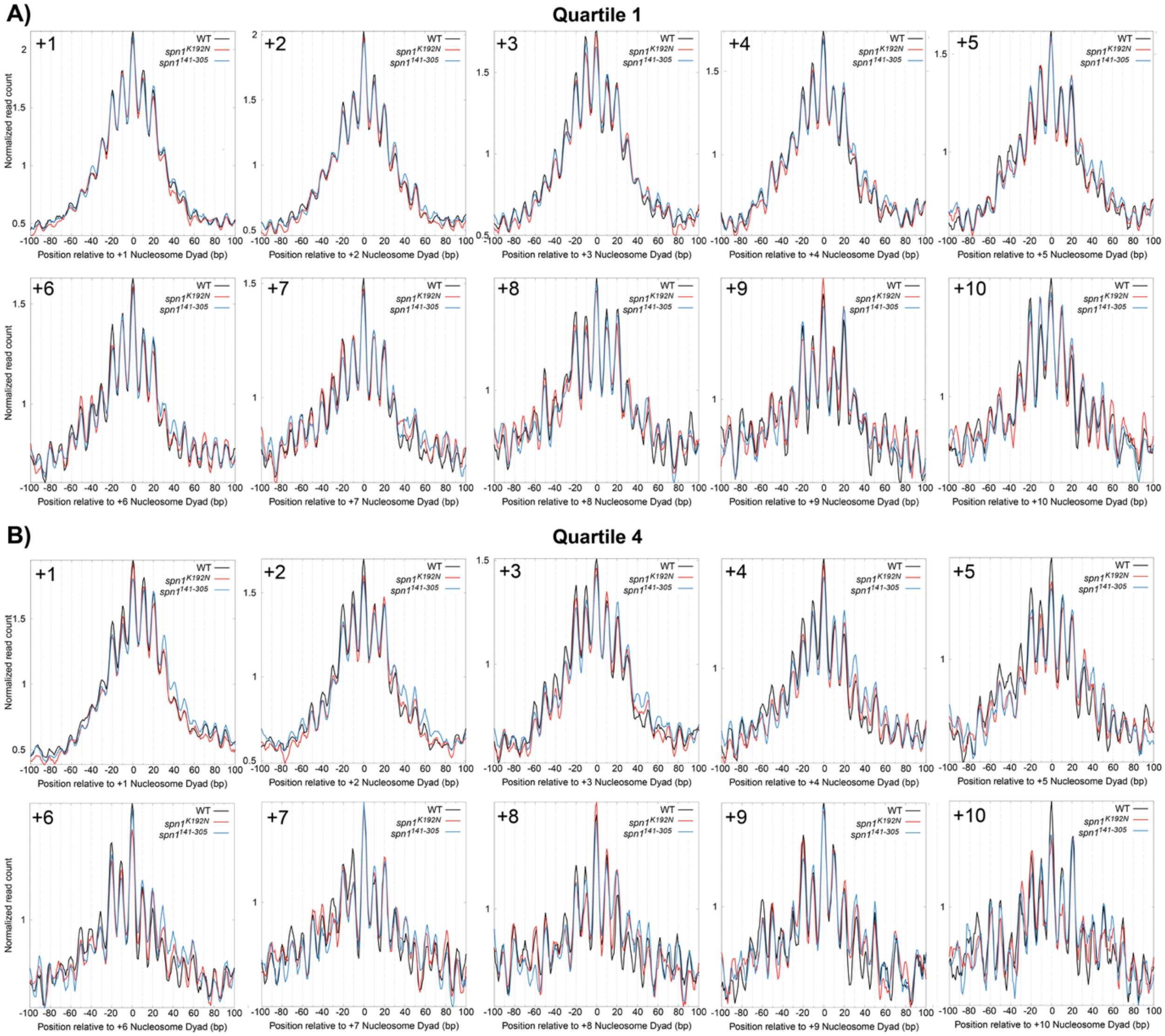

**Supplemental Figure S15: Subnucleosomal protections 102-112 bp in length in Spn1 mutants occupy the same rotational positions as wild-type cells, but with higher occupancy at rotational positions downstream of the dyad with a loss of occupancy at positions upstream of the dyad location.** Metagene plots of fragment midpoints 102 - 112bp in length mapped to each nucleosome dyad at the (A) lowest and (B) highest gene quartiles ranked by expression.

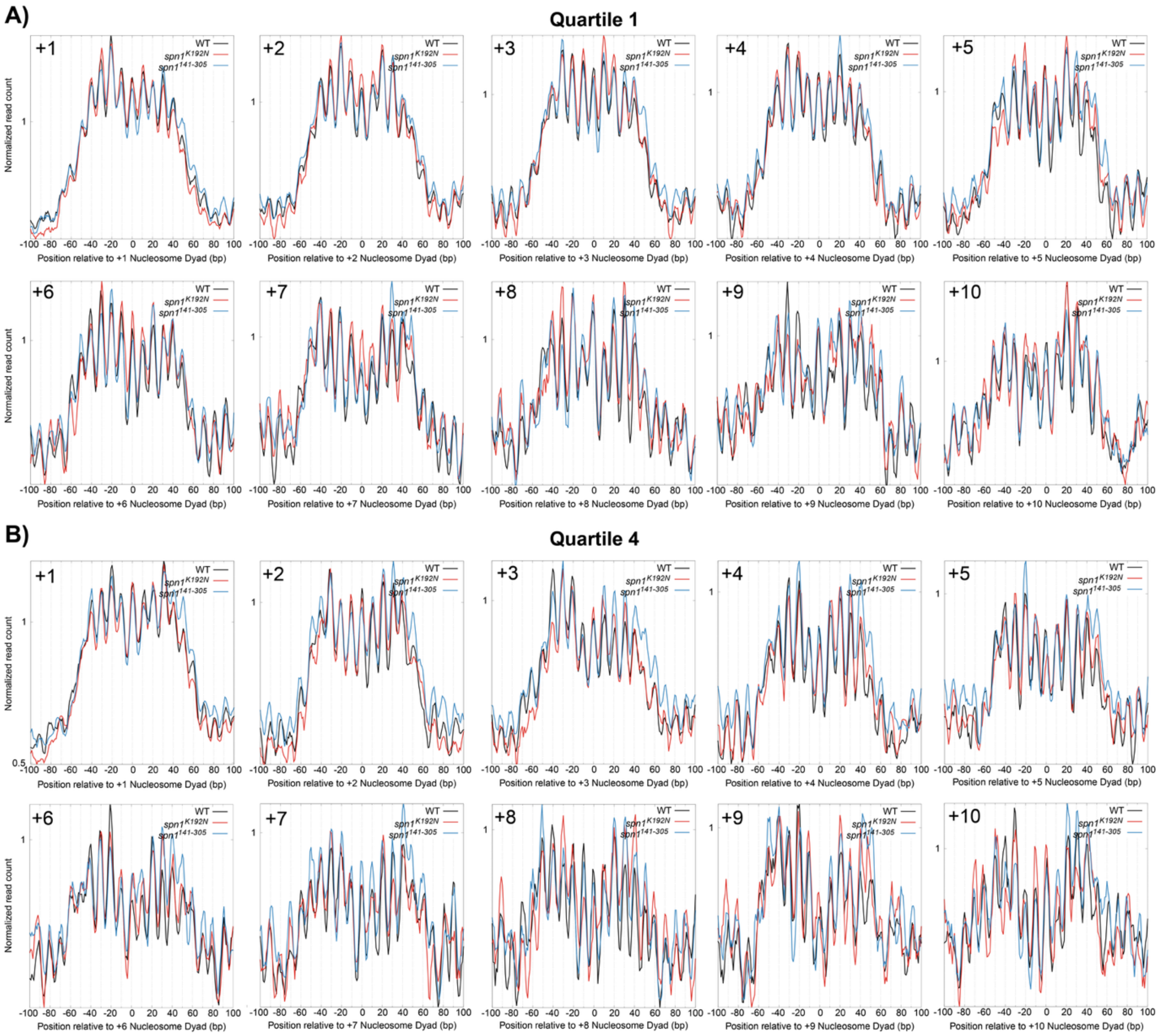


**Supplemental Figure S16: Subnucleosomal protections 61-71 bp in length in Spn1 mutants occupy the same rotational positions as wild-type cells, but with higher occupancy at rotational positions downstream of the dyad with a loss of occupancy at positions upstream of the dyad location in *spn1*$^{141\text{-}305}$.** Metagene plots of fragment midpoints 61 - 71bp in length mapped to each nucleosome dyad at the (A) lowest and (B) highest gene quartiles ranked by expression.

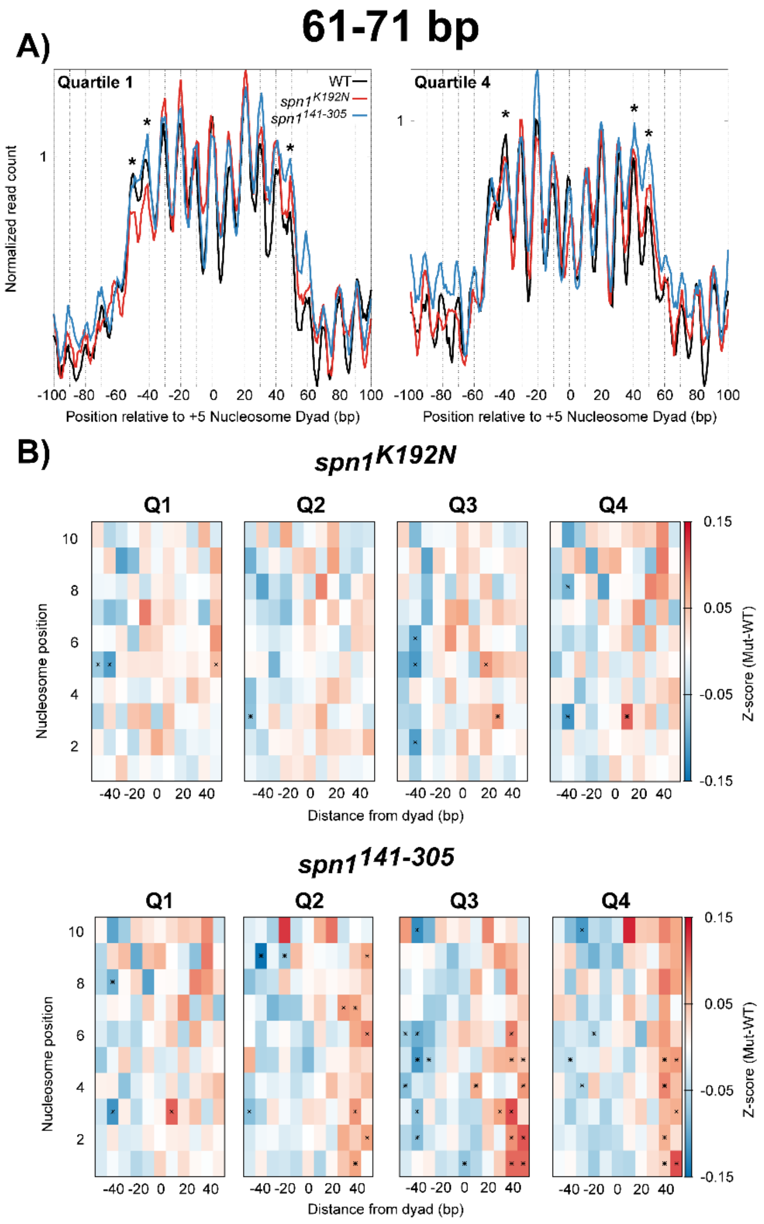


**Supplemental Figure S17:** Protected fragments 61-71 bp in length experience little shifts in Spn1 mutant expressing strains. (A) Metagene plots of fragment midpoints 61-71 bp in length mapped to the +5 nucleosome dyad at genes ranked by length. Asterisks indicate rotational positions with statistically significant change in Z-score. (B) Difference heatmaps between wild-type and mutant Spn1 expressing yeast strains scored by changes in 61-71 bp fragment occupancy at rotational positions surrounding each nucleosome dyad position at genes ranked by expression. Asterisks indicate rotational positions with statistically significant change in Z-score.

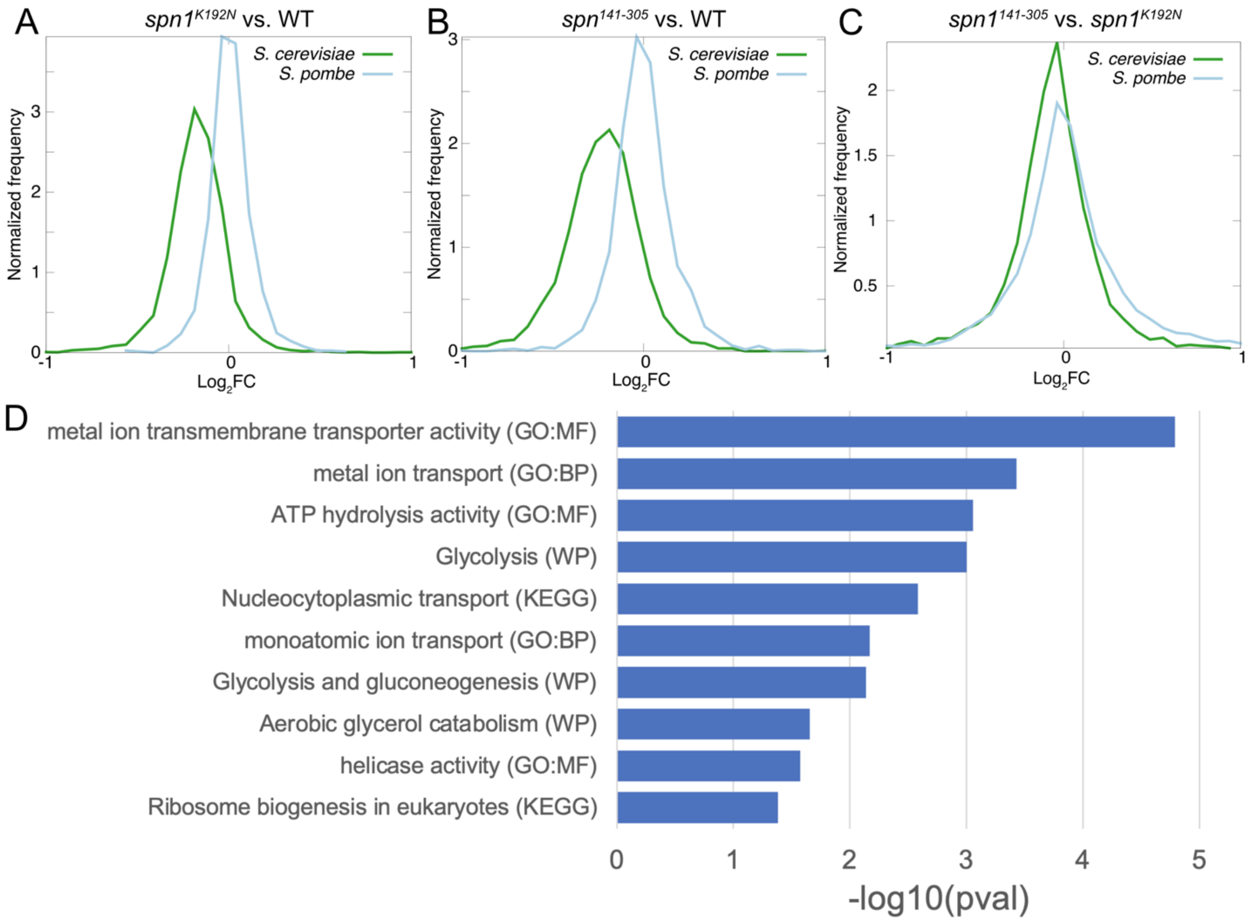


**Supplemental Figure S18: RNA-seq analysis of Spn1 mutants.** (**A**) Distribution of $log_2$(Fold Change) of *S. cerevisiae* and *S. pombe* genes for $spn1^{K192N}$ expressing yeast compared to wild-type. (**B**) Same as (**A**) for $spn1^{141\text{-}305}$ expressing yeast compared to wild-type. (**C**) Same as (**A**) for $spn1^{141\text{-}305}$ expressing yeast compared to $spn1^{K192N}$ expressing yeast. (**D**) Top 10 highest enriched terms obtained from gene set enrichment analysis of genes belonging to G1-G4. P-values adjusted for multiple testing correction are plotted on the x-axis. GO:MF denotes Gene Ontology: Molecular Function, GO:BP denotes Gene Ontology: Biological Process, WP denotes Wiki Pathways, KEGG denotes KEGG reactome.
